# Myocardial disarray drives metabolic inefficiency in human cardiomyocytes

**DOI:** 10.1101/2024.09.26.615170

**Authors:** Charlène Jouve, Andrea Ruiz-Velasco, Erminia Donnarumma, Rémi Le Borgne, Iman Momken, Céline Pereira, Magali Seguret, Eva Vermersch, Elodie Vimont, Ivan Nemazanyy, Jean-Marc Verbavatz, Luc Bertrand, Timothy Wai, Mathias Mericksay, Jean-Sébastien Hulot

## Abstract

Adult cardiomyocytes are embedded within a highly organized myocardial microenvironment that imposes critical geometric cues essential for the alignment and distribution of organelles and the shaping of their unique, rectangular cellular morphology. Despite the association of cardiomyocyte disarray with human heart disease, the functional consequences of this cellular disorganization remain poorly understood. Here, we leveraged micropatterned substrates to promote structural alignment in human induced pluripotent stem cell-derived cardiomyocytes (hiPSC-CMs), contrasting the effects of mechanical alignment on mitochondrial form and function with hiPSC-CMs cultured under standard unconstrained conditions.

Cardiomyocytes cultured under unconstrained conditions exhibited misaligned sarcomeres and a perinuclear mitochondrial distribution while micropatterned hiPSC-CMs developed linear myofibrils and reconfigured sarcomere and mitochondrial organization, which increased mitochondrial respiration without augmenting mitochondrial mass. Notably, micropatterned hiPSC-CMs exhibited an increased number of mitochondrial-associated membranes, as determined by proximity ligation assays and transmission electron microscopy, suggesting enhanced interactions between the sarcoplasmic reticulum and mitochondria.

Together, these findings demonstrate that mitochondrial-sarcoplasmic architecture and geometry are critical spatial features that ensure bioenergetic efficiency of cardiomyocytes. This work underscores the importance of cellular organization in cardiomyocyte metabolism and function, providing insights into the pathophysiology of cardiac diseases marked by cellular disarray.

## Introduction

The human cardiac muscle tissue presents a complex organization with a helical distribution of linear myocardial fibres which ensures its performance (1). The unique anisotropic architecture of the myocardium is not only essential to normal cardiac mechanics but also to other key functions, such as electrical signal propagation within the myocardial tissue (1, 2). This complex structure is formed during embryonic development and is supported by a highly structured microenvironment that imposes specific physical boundary conditions on cardiomyocytes. As a consequence, healthy cardiomyocytes are typically rod-shaped and aligned along a major elongation axis (3). This geometry configures the linear alignment of myofibrils and sarcomeres along the long axis of cells, ensures the intracellular organization of organelles, and the polarization of cardiomyocytes for proper cellular function (4, 5).

Many human heart diseases are associated with tissue and cell disorganization, leading to the disruption of the linear architecture of the myocardium. Myocardial disarray generally refers to disorganized cardiomyocyte spatial distribution, with loss of physiological fibre alignment and orientation at the histological level. Myocardial disarray was originally described as an archetypal feature of hypertrophic cardiomyopathy (HCM) (6, 7), but has also been found in other cardiac conditions including congenital heart disease, myocardial remodeling in ischemic heart diseases, genetic and acquired cardiomyopathies (8, 9).

In cardiomyocytes, myofibrils can also be disorganized, defining sarcomeric disarray. In patients with HCM, it was recently reported that myofibril disorganization was associated with mitochondrial dysfunction due to an improper organization around myofibrils (10). To ensure an efficient contraction of cardiomyocytes, mitochondria and the sarcoplasmic reticulum must be positioned in close vicinity of the contractile apparatus to provide the necessary ATP and calcium cycling required to power myofilament function. Genetic, infectious, and mechanical lesions that impinge upon cardiac and cardiomyocyte contractile function are associated with defect in the organization of these structural networks, however the causal and functional relationships between human cardiomyocyte architecture and function have yet to be elucidated. In part, this is because most human cellular cardiomyocytes models do not faithfully replicate the geometry observed in primary adult cardiomyocytes, which is correlated with differences in the observed bioenergetic profiles. To overcome this challenge, we developed a novel method for promoting sarcomere alignment and organization by culturing cardiomyocytes derived from human induced pluripotent stem cells (hiPSC-CMs) on bespoke micropatterned substrates composed of repeated lines of 30 µm-width separated with a cell repellent to impose specific geometric features. We then compare the alignment and organization of the micropatterned hiPSC-CMs to hiPSC-CMs cultured under standard unconstrained conditions.

## Results

### The culture of hiPSC-CMs as linear microfibers improves structural organization

We first generated ventricular hiPSC-CMs using the Wnt modulation protocol with glucose starvation (11, 12). The expression of pluripotency markers was validated by immunostaining and RT-qPCR (Figure S1A-C). Approximately 10 days after cardiac induction, cultures began displaying a contractile phenotype, and flow cytometry analysis on day 21 indicated an average proportion of 94.8 ± 6.0% TNNT2-positive cells (Figure S1D). The hiPSC-CM displayed typical expression of cardiac and sarcomeric proteins (Figure S1E-F). At 35 days of differentiation, the cells were enzymatically detached and replated onto micropatterned or unconstrained control (i.e. without linear micropatterning, referred to as non-patterned) coverslips coated with Matrigel (Figure 1A). Briefly, the micropatterned coverslips consist of repeated lines of 30 µm-width surrounded by a polymer with oxidation resistance and anti-adhesive properties (Figure 1B). On these micropatterned coverslips, Matrigel can only coat areas without the anti-adhesive agent thus creating boundaries for the generation of linear myofibrils as observed during embryogenesis. Otherwise, the culture condition is identical between micropatterned and control hiPSC-CMs.

**Figure 1.**
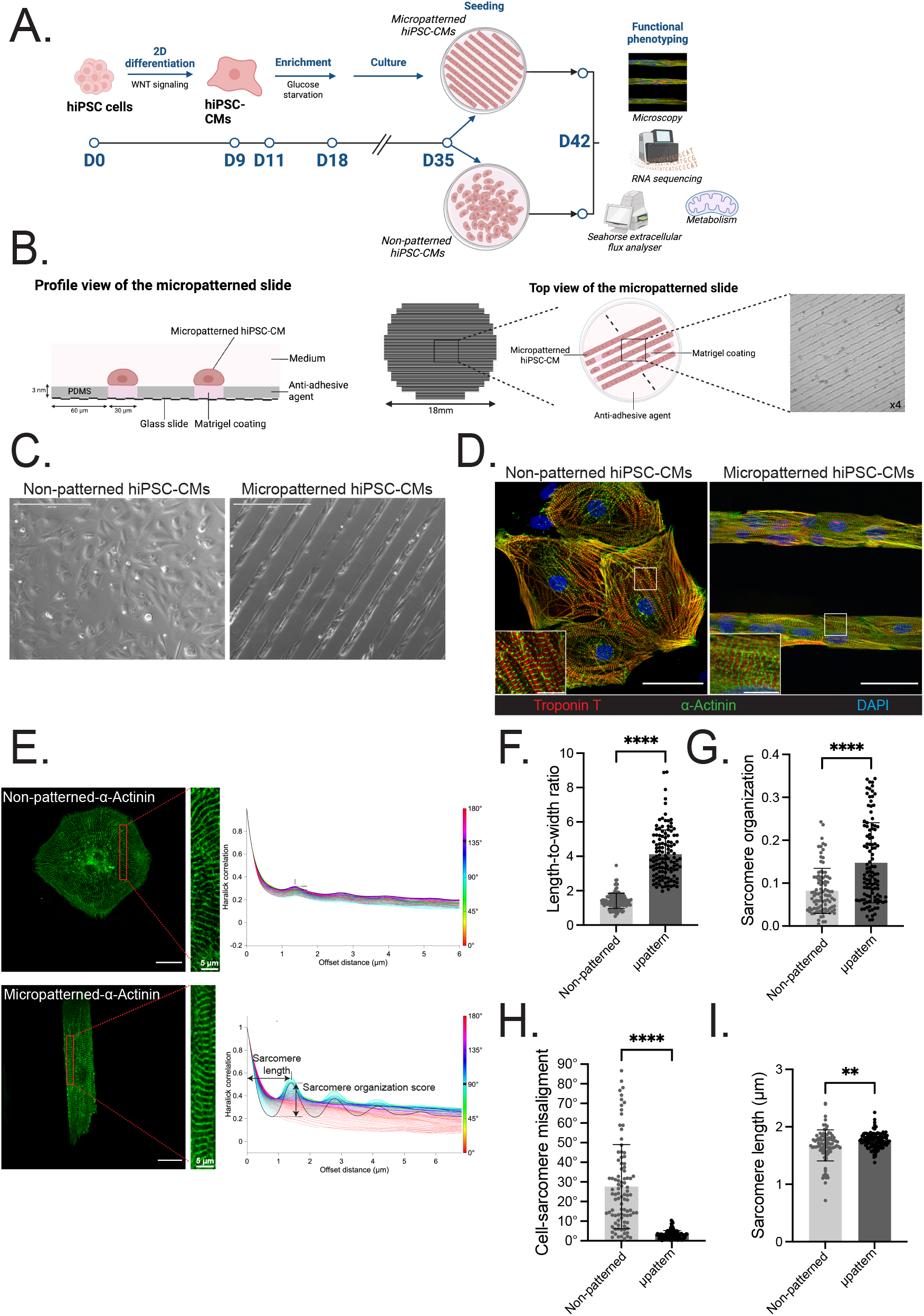
Linear micropatterning of human cardiomyocytes improves morphological features and sarcomere organization. **A**. Schematic overview of the study. On day 35 after differentiation initiation, hiPSC-CMs were seeded onto micropatterned or non-patterned slides for 7 days before functional phenotyping. **B**. Schematic representation of the profile (left) and top (right) view of the adhesive micropatterns. **C**. Representative phase-contrast images of non-patterned (left) and micropatterned (right) hiPSC-CMs, scale bar 400 µm. **D**. Representative immunofluorescence images of non-patterned and micropatterned hiPSC-CMs for *α*-sarcomeric actinin (green), cardiac Troponin T2 (red) at day 7 after seeding. DAPI (blue) was used to stain nuclei. The insert represents a zoomed view of sarcomeres. Global scale bar 100 µm, insert scale bar 10 µm. **E**. Automated measures of sarcomere organization and sarcomere length. (Left) Immunofluorescence images of non-patterned (top) and micropatterned (bottom) hiPSC-CMs. Scale bar 25 µm. (Right) Haralick correlation metric computed at multiple offset distances and angles to determine sarcomere organization and sarcomere length. Sarcomere organization score is the maximum amplitude of the decaying sinusoidal trace. Sarcomere length is the distance to the first peak. **F-I**. Scatter plots show quantification data for length-to-width ratio **(F)**, sarcomere organization **(G)**, cell-sarcomere misalignment **(H)**, and sarcomere length **(I)**. n = 138 non-patterned cells, n = 120 micropatterned cells; N = 3-4 differentiations. Results, mean *±* standard deviation. Statistical analyses: Unpaired t-test was used for F-I. ** p<0.01, **** p<0.0001 for the comparisons indicated.

Remarkably, the linear micropatterning substrate was able to spatially regulate the attachment and growth of hiPSC-CMs and promote multiple cardiomyocytes to form end-to-end connections, generating linear and aligned cardiac microfibers exhibiting a geometry comparable to that of primary adult cardiomyocytes (Figure 1C, S2A). In comparison, hiPSC-CMs cultured under standard unconstrained conditions showed a round to polygonal shape uncharacteristic of healthy, primary adult cardiomyocytes (Figure 1C). Seven days after replating, we assessed the cell morphology and sarcomere organization by immunofluorescence microscopy (Figure 1D). While all cells expressed the sarcomeric protein cardiac troponin T2 and the cardiac Z-disk protein *α*-actinin (Figure 1D, S2B), micropatterned hiPSC-CMs displayed an organized contractile machinery, characterized by an increase in the alignment of myofibrils throughout the cytoplasm of the cell and a better directionality within the microfiber. In contrast, hiPSC-CMs cultured under standard unconstrained conditions also showed sarcomere formation but with a lower degree of alignment and strikingly larger disorganization, consistent with previous reports (13) and characteristic of myocardial disarray.

To further characterize the organization of sarcomeres under both conditions, we used SarcOmere Texture Analysis method (14), a pixel-based image analysis for assessing sarcomere structure with *α*-actinin immunofluorescence (Figure 1E, S3A). We used a lower cell seeding density at D35 to facilitate cell segmentation and image analyses enabling single-cell quantification. The cell elongation (measured by the ratio of the major axis length to the minor axis length) was largely increased in micropatterned hiPSC-CMs (p<0.0001, Figure 1F), which reciprocally had a lower cell circularity index (Circularity = 4*π* x Area x Perimeter^−^2) (Figure S3B). Notably, micropatterned cardiomyocytes had a far greater number of aligned myofibrils, as evidenced by an increase in sarcomere score organization (p<0.0001, Figure 1G, S3A) and a decrease in the cell-sarcomere misalignment (p<0.0001, Figure 1H). Finally, we observed an increase in the resting sarcomere length in micropatterned hiPSC-CMs (p<0.01, Figure 1I).

Altogether, these data showcase the impact of micropatterning on the linear architectural organization and its ability to force sarcomere alignment in hiPSC-CMS.

### RNA-sequencing identifies changes in metabolic genes

We then asked whether the architectural changes could influence cardiomyocyte gene expression and performed mRNA-sequencing analysis. We first compared the native hiPSCs and the differentiated hiPSC-CMs by hierarchical clustering and tSNE analysis of their transcriptomes and confirmed a clear separation between their expression profiles (Figure S4A-C). Reactome gene set analysis (ReactomeGSA), a pathway analysis package, detected a down-regulation of pluripotency and proliferation, and an up-regulation of muscle formation and contraction in hiPSC-CMs, and confirmed a similar efficiency of the differentiation process in both micropatterned and non-patterned culture conditions (Figure S4D).

Despite marked structural differences between micropatterned and non-patterned hiPSC-CMs, the transcriptomic comparisons between micropatterned and non-patterned hiPSC-CMs showed no differential expression at the single-transcript level (Figure S4B). However, gene set analysis with ReactomeGSA detected 40 altered pathways involved primarily in metabolism and signal transduction (Figure 2A, S4E-F). Among these, four pathways indicated significant up-regulation of the lipid metabolism in micropatterned hiPSC-CMs (Figure 2B). More specifically, a significant up-regulation of genes involved in the metabolism of triglycerides and carnitines was observed (Figure 2C). Concordant with a change in metabolic pathways in micropatterned hiPSC-CMs, other up-regulated pathways including fatty acid oxidation, TCA, and respiratory electron transport pathways were observed (Figure 2C). On the other hand, genes associated with glycolysis were significantly down-regulated (Figure 2D).

**Figure 2.**
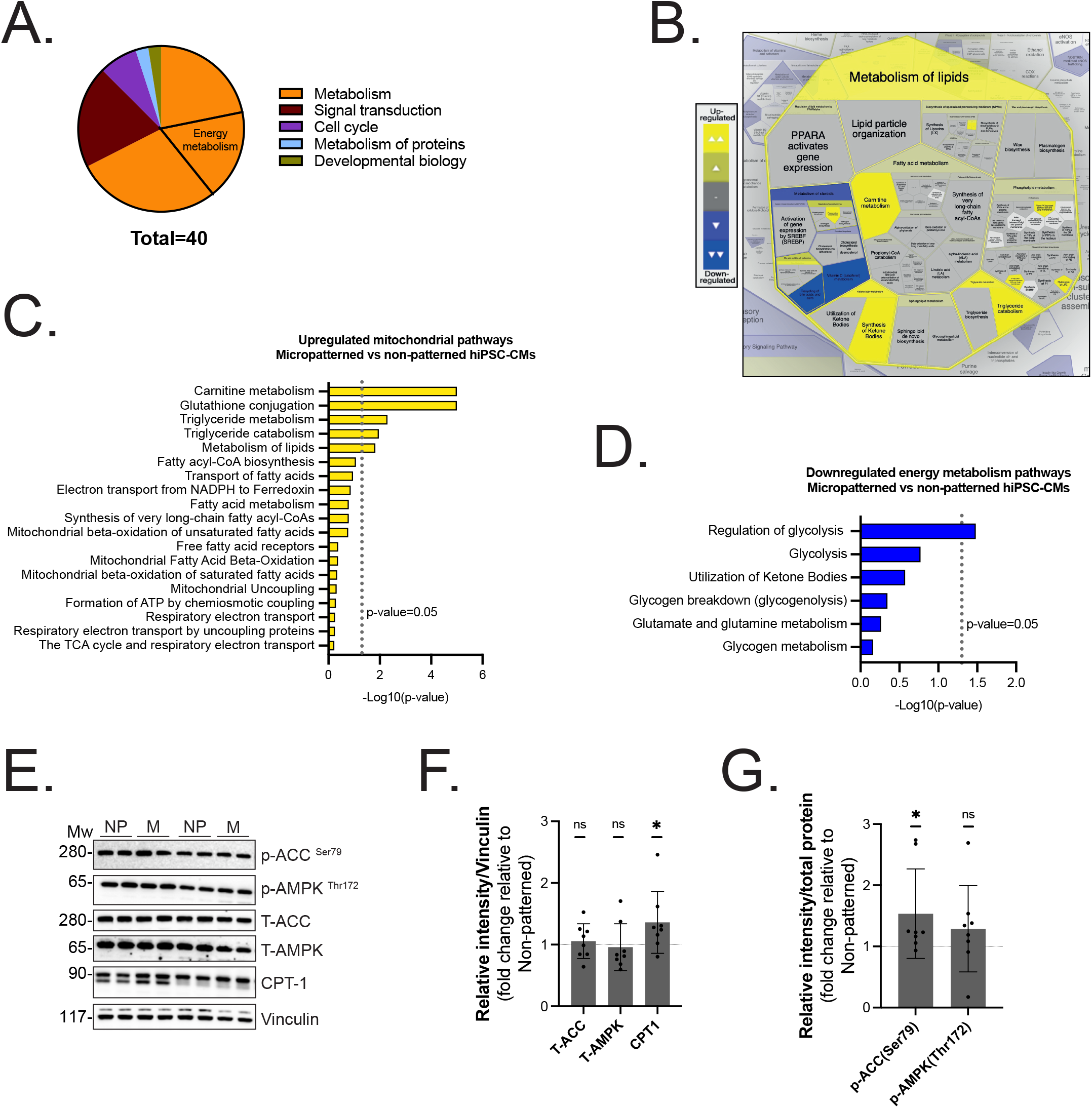
Transcriptomics and pathway analyses reveal changes in metabolic genes. **A**. Pie chart summarizing the classification of all the significantly changed pathways (FDR<0.01) in micropatterned vs. non-patterned hiPSC-CMs as found with ReactomeGSA. **B**. Voronoi diagram displaying the significantly altered lipid metabolism pathways (FDR<0.01) in micropatterned vs. non-patterned hiPSC-CMs (ReactomeGSA). The upregulated pathways are colored yellow, while the downregulated pathways are colored blue. **C**. Upregulated energy metabolism pathways identified by ReactomeGSA comparing micropatterned vs. non-patterned hiPSC-CMS showing their corresponding p-value as -Log10(p-value). **D**. Downregulated energy metabolism pathways identified by ReactomeGSA comparing micropatterned vs. non-patterned hiPSC-CMS showing their corresponding p-value as -Log10(p-value). **E**. Representative Western blot images showing the expression of ACC (total and phosphorylated at Ser79), AMPK (total and phosphorylated at Thr172), and CPT1 on non-patterned (NP) and micropatterned (M) hiPSC-CMs. Vinculin was used as loading control. **F**. Quantification of total proteins ACC, AMPK, and CPT1 in micropatterned hiPSC-CMs (n = 8, expressed as fold change relative to non-patterned for each cardiac differentiation). **G**. Quantification of p-ACC^*Ser*79^/ACC and p-AMPK^*T hr*172^/AMPK ratio in micropatterned hiPSC-CMs (n = 8, expressed as fold change relative to non-patterned for each cardiac differentiation). Results, mean *±* standard deviation. Statistical analyses: Wilcoxon signed-rank test was used for F, and G. *p<0.05, ns: not significant for the indicated comparisons.

Given that cardiac energy metabolism relies primarily on fatty acid oxidation to fuel the bioenergetic needs of cardiomyocytes, we further focused on the carnitine pathway (Supplementary Table1) and studied the protein expression of three major components of the pathway, including Acetyl-CoA carboxylase (ACC), AMP-activated protein kinase (AMPK), and the carnitine palmitolytransferase (CPT1). We found that while total ACC and AMPK steady were not different between conditions, the levels of the fatty acid mitochondrial transporter CPT1 were increased in micropatterned vs. non-patterned hiPSC-CMs (Figure 2E-F). In line, we found a significant increase in the pACCSer79:ACCtotal in micropatterned hiPSC-CMs (Figure 2G), suggesting a lower conversion of acetyl-CoA to malonyl-CoA which is a known inhibitor of fatty acid transport into mitochondria. Altogether, these changes point to a shift in substrate preference in the micropatterned hiPSC-CMs and support that the architectural organization of hiPSC-CMs drives their metabolic rewiring, promoting the use of fatty acid sources by mitochondria.

### Changes in respiratory functions according to sarcomere organization

As our RNA-sequencing data indicated a significant change in pathways related to energy metabolism, we tested whether the architectural manipulation led to changes in respiratory functions. We characterized mitochondrial respiration by measuring oxygen consumption rate (OCR) in micropatterned vs. non-patterned hiPSC-CMs in basal conditions and after the addition of oligomycin (an inhibitor of ATP synthase), FCCP (a mitochondrial electron uncoupler), rotenone and antimycin A (complexes I and III inhibitors) (Figure 3A). We found that micropatterned hiPSC-CMs displayed a strong increase in OCRs associated with basal respiration, ATP production (couples respiration), and maximal respiration as compared to non-patterned hiPSC-CMs (Figure 3A-B). We observed no changes in spare respiratory capacity and proton leak, suggesting that the observed overall increase in oxidative phosphorylation (OXPHOS) capacity is not the result of uncoupling (Figure S5A). We observed no differences in the ECAR values between micropatterned-CMs and non-patterned-CMs in response to the addition of glucose, indicating that mitochondrial utilization of pyruvate derived via glycolysis is equally efficient in both cell types. Similarly, the subsequent addition of oligomycin yielded a commensurate and comparable increase in ECAR, demonstrating that the inhibitory effect on ATPase activity is equivalent (Figure 3C-D, S5B). When comparing the energy phenotype from micropatterned-CMs and non-patterned-CMs, we observed that micropatterned cells adopted a more energetic profile due to an increased capacity to consume oxygen, whereas non-patterned cells were more glycolytic (Figure 3E).

**Figure 3.**
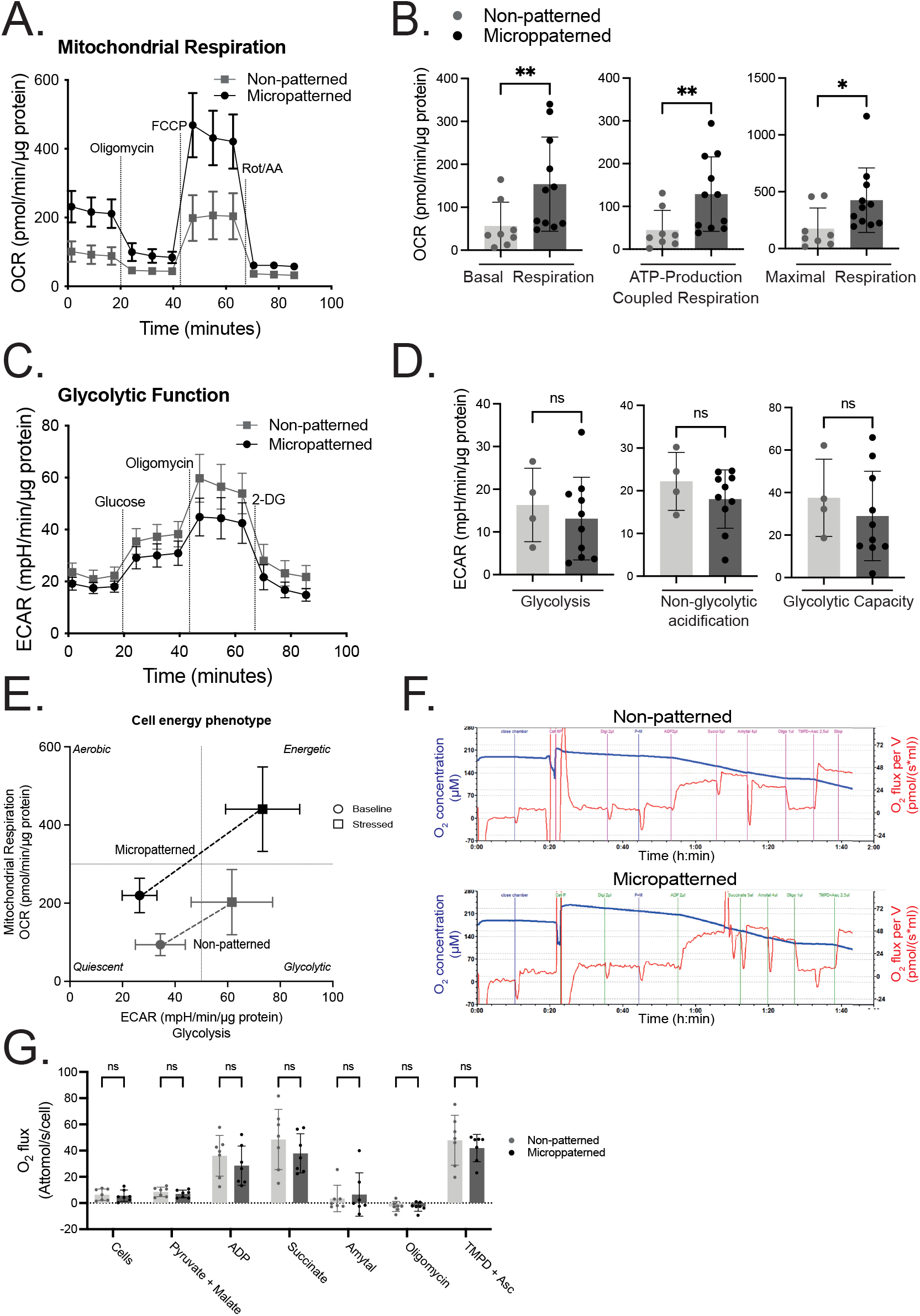
Mitochondrial respiration is increased in micropatterned hiPSC-CMs. **A**. Real-time oxygen consumption rate (OCR) measurements of micropatterned and non-patterned hiPSC-CMs by Seahorse extracellular flux analyzer. Cells were treated with the ATP synthase inhibitor oligomycin, the respiratory uncoupler FCCP, and the respiratory chain blockers rotenone and antimycin A. **B**. Scatter plots show basal respiration, production of ATP, and maximal respiration. N = 8-11 differentiations. **C**. Real-time extracellular acidification rate (ECAR) measurements of hiPSC-CMs non-patterned and micropatterned by Seahorse extracellular flux analyzer. Cells were treated with glucose, ATP synthase inhibitor oligomycin, and glucose analog 2-DG. **D**. Scatter plots show glycolysis, non-glycolytic acidification, and glycolytic capacity. N = 4-12 differentiations. **E**. Cell energy diagram showing normalized oxygen consumption rate (OCR) and normalized extracellular acidification rate (ECAR) of micropatterned and non-patterned hiPSC-CMS at the basal (circle points) and stressed (square points) conditions. **F**. Representative profiles of oxygen consumption of non-patterned (top) and micropatterned (bottom) hiPSC-CMs measured in permeabilized cells with complex protocol using Oxygraph-2k high resolution respirometry. Digitonin (Dig, 10 µg/ml), pyruvate (P, 1 mM) and malate (M, 4 mM), ADP (2 mM), succinate (10 mM), amytal (2 mM), oligomycin (10 µM), and TMPD (0.5 mM) combined with ascorbate (2 mM) were added sequentially. Blue line stands for decreasing oxygen concentration in the oxygraphy chamber containing permeabilized cells. Red line stands for the changes of oxygen flux per volume. **G**. Oxygen flux (attomol/sec/cell) quantified in non-patterned and micropatterned hiPSC-CMs. n = 7 in each group from 3 independent differentiation. Results, mean *±* standard deviation. Statistical analyses: Mann-Whitney was used for B, and D, two-way ANOVA was used for G. * p<0.05, ** p<0.01, ns: not significant for the indicated comparisons.

To determine whether the intrinsic organization of mitochondria within patterned cardiomyocytes is responsible for the increased OXPHOS capacity observed in micropatterned cells, we measured oxygen consumption rates by high-resolution respirometry in isolated mitochondria using the Oroboros system (Figure 3F-G). Isolated mitochondria from micropatterned and non-patterned cells showed similar oxygen flux rates in the presence of pyruvate, malate and ADP (state 3) and succinate. As in intact cells, we observed no differences in state 4 respiration (addition of oligomycin) indicating that mitochondria are equally coupled. Finally, the activity of cytochrome *c* oxidase, the terminal enzyme of the respiratory chain that consumes oxygen in mitochondria, was measured by injecting TMPD and Ascorbate, revealing indistinguishable levels of activity.

Taken together, our data indicate that the linear architectural organization of sarcomeres drives increase bioenergetic efficiency by creating an intracellular environment that can maximize mitochondrial respiration without altering the expression of the components of the OXPHOS system. In the absence of structural remodeling of hiPSC-CMs, cells retain a more glycolytic preference due to an inability to boos OXPHOS activity.

### Effect of linear micropatterning on mitochondria organization

To explore the cellular basis of increased OXPHOS capacity of mitochondria, we used MitoTracker™ dye as an indicator to assess the mitochondrial network morphology from micropatterned and non-patterned hiPSC-CMs. As shown in Figure 4A, the mitochondrial network from micropatterned hiPSC-CMs appeared to be more homogeneously distributed and organized along the myofibrils, compared to the non-patterned hiPSC-CMs where mitochondria accumulated around the nucleus (Figure S6A). This was evidenced by an increase in mitochondrial directionality within the microfiber (Figure 4B). IMARIS analysis revealed that global mitochondrial volume was increased in the micropatterned condition compared to the non-patterned condition albeit modestly (1.065%) (Figure 4C-D), without a significant change in cardiomyocyte cell surface (Figure S6D). The distribution of mitochondrial volume was identical to the non-patterned condition (Figure S6E). No difference was found in the number of mitochondria by imaging analysis (Figure 4E) or by quantification of mtDNA content (Figure 4F). Mitochondrial Orange signal labeling actively respiring cells (Figure 6B) and Mitotracker mitochondrial mass (Figure 6C) were similar in both conditions.

**Figure 4.**
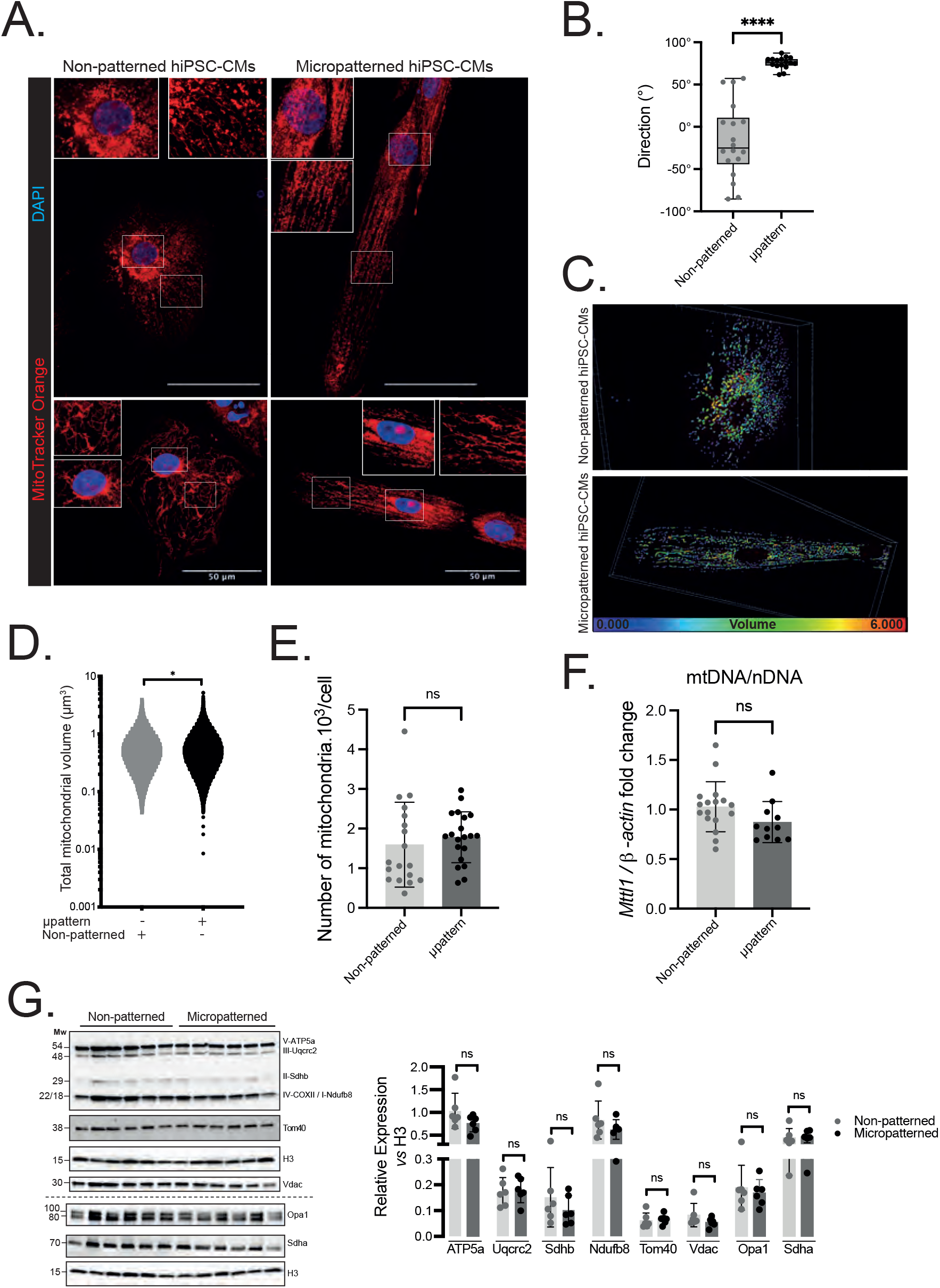
Micropatterning impacts mitochondrial distribution independently of mitochondrial mass. **A**. Mitochondrial distribution stained by MitoTracker™ Orange (red) and DAPI (blue) and analysed by confocal microscopy. The insert represents a zoomed view of mitochondrial network. Global scale bar 50 µm, insert scale bar 10 µm. **B**. Quantitative analysis of mitochondrial direction measured with Fiji software. n = 18 non-patterned cells, n = 16 micropatterned cells; N = 2 differentiations. **C**. Representative pictures obtained with MitoTracker™ Orange and IMARIS software. Each mitochondrion is identified by a color in accordance with its volume (the smallest mitochondria are blue and the largest are red). **D**. Total mitochondrial volume quantified by IMARIS. n = 28,675 non-patterned cells, n = 64,919 micropatterned cells; N = 3 differentiations.**E**. Scatter plots show the number of mitochondria per cell. n = 18 non-patterned cells, n = 20 micropatterned cells; N = 3 differentiations. **F**. Mitochondrial DNA (mtDNA) content in non-patterned (n = 17) and micropatterned (n = 11) hiPSC-CMs, quantified by amplification of the mitochondrial *Mttl1* gene relative to nuclear gene *b-Actin*. N = 3 differentiations. **G**. Cardiac levels of ATP5A, UQCRC2, SDHB, I-NDUFB8, TOMM40, VDAC, OPA1, and SDHA were analysed by western blot in non-patterned and micropatterned hiPSC-CMs. Total Histone3 was used as loading control (n = 6). Results, mean *±* standard deviation. Statistical analyses: Unpaired t-test was used for C, Mann-Whitney was used for D through G. * p<0.05, ns: not significant for the indicated comparisons.

Concordant with the Seahorse and Oroboros respiration measurements (Figure 3) and in line with the RNA-sequencing data, the steady state levels of OXPHOS proteins and subunits of mitochondrial complexes required for mitochondrial biogenesis (ATP5A, UQCRC2, SDHB, I-NDUFB8, TOMM40, VDAC, OPA1, and SDHA) were not significantly different between micropatterned and non-patterned hiPSC-CMs (Figure 4G). Taken together, our data indicate that the increased OXPHOS capacity driven by structural reorganization of micropatterned hiPSC-CMs is not the result of increased mitochondrial biogenesis nor in changes in mitochondrial content.

### Increased interactions between sarcoplasmic reticulum and mitochondria following linear organization of sarcomeres

To gain insights in the mechanism underlying increased bioenergetic output in micropatterned cardiomyocytes, we turned our attention on the spatial organization and interaction of myofilaments, mitochondria, and sarcoplasmic reticulum (SR). In line with the immunostaining results, transmission electron microscopy (TEM) analyses undercover an ultrastructural improvement in micropatterned hiPSC-CMs with an organization of the mitochondrial network relative to the myofibrils and the SR closer to the one observerd in primary adult cardiomyocytes (Figure5A, S7A). As the SR interacts physically and functionally with mitochondria through mitochondria-associated membranes (MAMs), we tested proteins that tether the two organelles together (Figure 5B). IP_3_R1 on the SR interacts with VDAC1 on the outer mitochondrial membrane through the molecular chaperone glucose-regulated protein 75 (GRP75) (15). We first checked that the expression levels of IP_3_R1, VDAC1, and GRP75 proteins were not significantly different between the two conditions (Figure 5C). We then assessed whether the interconnection between SR and mitochondria is modified and whether it may be linked to the increased mitochondrial oxidative function in linear architectural pattern. We evaluated the contact point between the two organelles using the *in situ* PLA assay (Figure S7B) as previously described (16, 17). The IP_3_R1-VDAC1, IP_3_R1-GRP75, and VDAC1-GRP75 interactions were significantly increased in micropatterned hiPSC-CMs, indicating that the physical SR-mitochondria interconnections are enhanced (Figure 5D).

**Figure 5.**
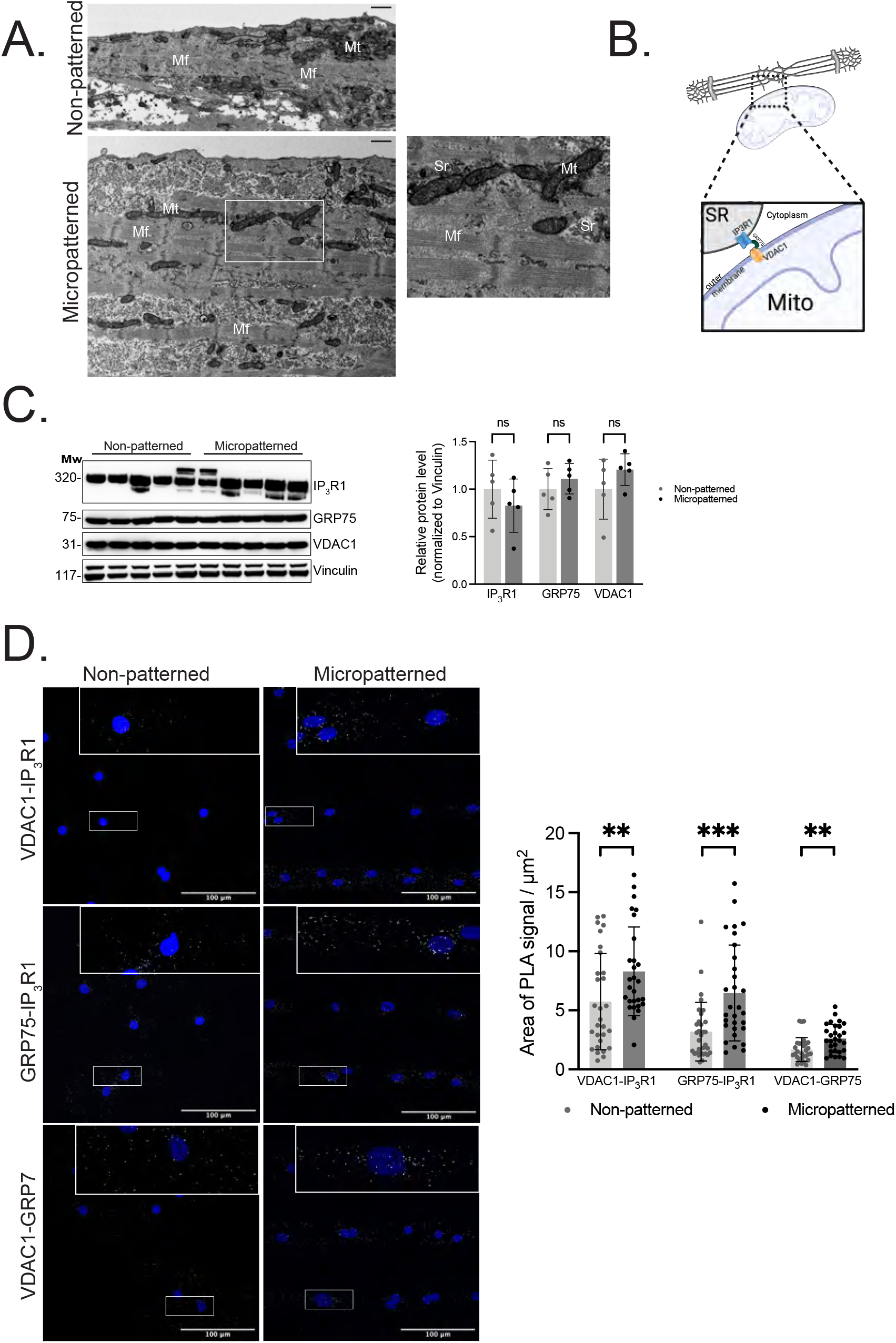
Increase of mitochondria-SR contact sites by structural reorganization with micropatterns. **A**. Transmission electron microscopy (TEM) images of the cellular structure of non-patterned (top) and micropatterned (bottom) hiPSC-CMS in transverse cross-sections; scale=1 µm. Mf indicates myofibril; Mt mitochondria; and SR sarcoplasmic reticulum. **B**. Schematic representation of the VDAC1/GRP75/IP_3_R1 complex at MAM interface. **C**. Western blot and quantitative analysis of VDAC1, GRP75, and IP_3_R1 protein abundance in non-micropatterned and micropatterned hiPSC-CMs. n = 5 differentiations / conditions. Vinculin was used as a loading control. **D**. Left: representative PLA images. The insert represents a zoomed view of PLA signal. Each white dot corresponds to a positive PLA signal. Right: quantitative analysis of VDAC1-IP_3_R1, and GRP75-IP_3_R1, and VDAC1-GRP75 interactions signal in non-patterned and micropatterned hiPSC-CMs. n = 10 images / conditions, N = 3 differentiations. Scale bar 100 µm. Results, mean *±* standard deviation. Statistical analyses: Mann-Whitney test was used for C and D. ** p<0.01, *** p<0.001, ns: not significant for the indicated comparisons.

These results demonstrate that linear architectural reorganization leads to an increase in the SR-mitochondria interactions.

## Discussion

Many cardiac diseases are associated with remodeling processes that profoundly alter the architecture of myocardial tissue (18). Myocardial disarray was initially observed at the histological level (8), but the disorganization also occurs at the cellular level. The spatial organization of the sarcomere is known to be critical for proper tension generation during myocyte excitation. However, the relationships between architectural changes in inter-organelle interactions and their functional consequences during heart failure in humans are poorly understood. This is partly due to the lack of human *in vitro* models that reproduce cellular disarray that underscores myocardial dysfunction in patients. Here, we used a micropatterning strategy to induce architectural organization and sarcomere alignment as observed in normally formed human cardiomyocytes. Strikingly, we found that enforcing linear architectural organization in human cardiomyocytes enhances the bioenergetic efficiency of mitochondria. Our data indicate that the improved mitochondrial respiration following linear architectural organization is not explained by an increase in the mitochondrial mass or biogenesis but rather by a cellular organization of the mitochondrial network relative to the myofibrils that recapitulate the organization observed in isolated mammalian primary cardiomyocytes with mitochondria in micropatterned hiPSC-CMs that were mainly located in intrafibrillar regions of the cardiomyocyte. Conversely, reduced mitochondrial respiration in disorganized cardiomyocytes was not associated with a reduced number of mitochondria or abnormal expression of mitochondrial proteins, highlighting the importance of the quality and organization of mitochondria in facilitating optimal respiratory capacity.

Mitochondria are symbiotic partners with the other cellular components. Cross-talk between the sarcoplasmic reticulum (SR) and mitochondria is essential for several cellular processes, including lipid synthesis and transport (19), mitochondrial function (20) and regulation of Ca^2+^ homeostasis (21). The SR physically and functionally interacts with mitochondria through mitochondria-associated membranes (MAMs). MAMs are microdomains that allow efficient transfer of calcium and lipids between the SR and mitochondria, further regulating metabolism (19, 22). These interactions are mediated by tethering proteins, including the voltage-dependent ion channel (VDAC1) on the outer mitochondrial membrane, the inositol 1,4,5-triphosphate receptor (IP_3_R)-1 complex on the SR, and the molecular chaperone glucose-regulated protein 75 (GRP75) (15). While the expression levels of IP_3_R1, VDAC1, and GRP75 proteins were not affected by the architectural constraints, in situ proximity ligation assay showed that the integrity of MAMs was disrupted in disorganized non-patterned hiPSC-CMs, indicating a reduction in the number of SR-mitochondria contact sites. The importance of MAMs in the pathogenesis of heart failure is increasingly recognized, but the mechanisms regulating MAM formation and their function in cardiomyocytes remain elusive. Our findings highlight the key role of the anisotropic structure of the myocardium in the correct formation of MAMs. Whether our micropatterning approach would further help to understand the role of MAMs in failing hearts or in cardiomyopathies (e.g., using mutant hiPSC-CMs) deserves further investigation.

Our results also position mitochondrial bioenergetic inefficiency as a primary maladaptive event that occurs in response to cellular dysmorphology of cardiomyocytes, which helps reconcile the long-standing yet unexplained observation of cardiomyocyte dysmorphology and mitochondrial dysfunction that manifests in preclinical models of heart failure and human patient alike (23–26). Previous animal studies have demonstrated that a cardiomyocyte-specific ablation of mitochondrial proteins that control mitochondrial energy metabolism, mitochondrial dynamics, or inter-organellar contacts with mitochondria trigger heart failure characterized by cardiomyocyte dysmorphology and dysfunction (26–31). A transgenic mouse model with a cardiac-specific ablation of the mitochondrial peptidase YME1L showed mitochondrial fragmentation and progressive development of dilated cardiomyopathy, heart failure, and reduced lifespan (32). Similarly, mitochondrial complex I-deficient animals showed impaired systolic function and reduced cardiac output (33). These mouse studies indicated that mitochondrial dysfunction in heart failure is not only a consequence but rather a contributor to cardiac dysfunction. While mutations in mitochondrial genes can cause heart failure in a subset of patients with mitochondrial disease, they account for only a small portion of all inherited cardiomyopathies and of heart failure in humans (34, 35). Conversely, mitochondrial dysfunction is observed in cardiomyocytes from hearts with common cause of heart failure. Our data support the notion that altering cellular morphology and structural organization of human cardiomyocytes is sufficient to modulate mitochondrial energy metabolism, which helps reconcile the discrepancy between the abundance of mitochondrial dysfunction observed in idiopathic and non-mitochondrial inherited cardiac diseases and the paucity of primary mitochondrial mutations that result in cardiac dysfunctions in humans. This would yield a model in which genetic, infectious, or chemical lesions that primarily disrupt cardiomyocyte architecture would lead to defects in mitochondrial bioenergetics. Our findings add to this concept by showing a sequence in which changes in mechanical constraints primarily lead to disorganization of the mitochondrial network and impaired mitochondrial oxidation.

Finally, a metabolic switch away from fatty acid oxidation and towards glucose utilization is often observed in the failing heart ((36, 37). Here, we found that organized micropatterned hiPSC-CMs had a higher metabolic preference for oxidation over glycolysis compared with non-patterned hiPSC-CMs, which had proportionally higher use of glycolysis. This observation was associated with a shift in substrate preference for lipids and an environment favoring the transfer of fatty acids to mitochondria, supporting that architectural organization leads to an improved mitochondrial function that is more dependent on fatty acid sources. This could be supported by the increased integrity of MAMs, which are also known to facilitate lipid trafficking between the two organelles (38). In addition, MAMs are also enriched in enzymes that mediate fatty acid metabolism as well as their ligation to coenzyme A (38, 39). Taken together our results indicate that architectural disorganization of the mitochondrial network in cardiomyocytes primarily leads to disruption of the MAMs, reduced lipid metabolism, and lower mitochondrial respiration. Consistently, nutritional intervention with a high-fat diet in transgenic mice with the mutations associated with mitochondrial cardiomyopathies resulted in improved substrate utilization and regression of the dilated cardiomyopathy phenotype, while mitochondrial fragmentation was still observed (32, 40).

Our experimental model was designed to specifically study the role of geometric constraints on cardiomyocyte biology, which may be a limitation in recapitulating the complexity of pathological stimuli that occur during heart failure. Mitochondrial function is dependent on other stress factors such as ER stress, oxidative stress, or cell apoptosis. Future experiments combining our micropatterning approach with additional stressors will provide further insight into this issue.

## Conclusions

Architecture drives the efficiency of mitochondrial respiration in cardiomyocytes through a more efficient mitochondrial/sarcoplasmic structural organization and an increased mobilization of fatty acid release into the mitochondria for oxidation.

## Materials and methods

### hiPSC culture and maintenance

The human induced pluripotent stem cell SKiPSC-31.3 line from a healthy male adult volunteer, has been used here and was previously reported (41–44). HiPSCs were seeded on stem cell-qualified Matrigel-coated (Corning, Corning®, NY, USA) plates and maintained in mTeSR™ Plus medium (STEMCell Technologies) in a humidified incubator (5% CO_2_, 37°C). When reaching 70 to 80% confluency, cells were passaged in clumps by scraping with a pipette tip. Pluripotency was assessed by immunostaining and PCR with reverse transcription (RT-qPCR) (Figures S1) and routinely tested for mycoplasma. Genomic stability was assessed by the detection of recurrent genetic abnormalities using the iCS-digital™ PSC test, provided as a service by Stem Genomics (https://www.stemgenomics.com/), as described previously (45).

### Cardiac differentiation

When hiPSCs reached a 90% confluency, differentiation was carried out according to a protocol adapted from (46). Briefly, mTeSR™ Plus was re-placed with RPMI-1640 supplemented with B27 supplement minus insulin (Thermo Fisher Scientific, Waltham, MA, USA) and 6 µM CHIR-99021 (Abcam) and maintained in a 5% CO_2_/air environment for 48 hours. The medium was then changed to RPMI-1640/B27 minus insulin for 24 hours and switched to RPMI-1640/B27 minus insulin supplemented with 5 µM IWR-1 (Sigma-Aldrich, St. Quentin Fallavier, France) for 48 hours. On day 5, the medium was changed back to basal medium for 48 hours. From day 7 onwards, cells were placed in RPMI-1640 with B27 supplement plus insulin (Thermo Fisher Scientific). On day 11, beating hiPS-CMs were subjected to glucose starvation in RPMI-B27 without glucose (Thermo Fisher Scientific) for 3 days. Cells were then dissociated using 0.05% trypsin (Thermo Fisher Scientific) for 10 minutes and seeded at 0.3 × 10^6^ cells/cm^2^ in RPMI-1640/B27 plus insulin for 24 hours. Then the cells were subjected to a second round of glucose starvation for 3 days. These two rounds of glucose starvation have been shown to increase dramatically the percentage of cardiomyocytes obtained (11). By day 18, cells were cultured in RPMI-1640/B27 plus insulin and the medium was changed every two days.

### Growing cells on micropatterned surface

18 mm round micropatterned coverslips were purchased from 4Dcell (Montreuil, France), with custom shape (repeated lines of 30 µm-width separated with a cell repellent). On day 35 hiPSC-CMs were seeded into Matrigel-coated micropatterned coverslips at a density of approximately 62.5-125 × 10^3^ cells/cm^2^ and allowed to grow for 7 to 10 days with RPMI-1640/B27 plus insulin at 37°C in 5% CO2 atmosphere. hiPSC-CMs cultured under standard unconstrained conditions were used as control (non-patterned condition).

### Flow Cytometry

hiPSC-CMs were dissociated into single cells with 0.05% trypsin-EDTA and stained with the fixable Zombie NIR™ Viability Kit (BioLegend®) according to the manufacturer’s protocol. Then, collected cell pellets were fixed and permeabilized using Inside Stain kit (Miltenyi Biotech) for 10 minutes at room temperature. Cells were incubated with fluorescent labelled antibodies (Supplementary Table2) for 10 min at room temperature. Analysis was performed on BD Biosciences FACS LSR Fortessa X-20 instrument with at least 30,000 cells. Results were processed and plotted using FlowJo v10 software (FlowJo, LLC).

For quantification of mitochondrial content, mitochondria of dissociated non-patterned and micropatterned hiPSC-CMs (day 42) were stained using 200 nM MitoTracker™ Green FM (Thermo Fisher Scientific). Live cells were incubated with the dyes in Ca^2+^/Mg^2+^-rich HBSS for 30 minutes at 37°C and 5% CO2, followed by a PBS wash. Acquisition processing and analysis were performed as described above.

### Immunostaining and imaging

hiPSC-CMs were cultured on Matrigel-coated micropatterned or standard unconstrained coverslips. After one week, cells were sequentially fixed with 4% paraformaldehyde (Electron Microscopy Sciences) for 10 min and then permeabilized and blocked with 0.5% Triton X-100, 2% bovine serum albumin in PBS (blocking solution) for 1 hour. Subsequently, primary antibody incubation was performed overnight at 4°C in 1:10 diluted blocking solution (Supplementary Table2). Subsequently, the cells were incubated with Alexa Fluor 488 or 546-conjugated goat anti-mouse or goat anti-rabbit antibodies (Supplementary Table2) for 1 hour at room temperature and counter-stained with DAPI nuclear dye (1:10,000). Coverslips were mounted with Dako Faramount Aqueous Mounting (Agilent) and dried for 24 hours. Fluorescent images were captured on a Cam mounted on a Leica TSC SP8 microscope. Image processing and analysis were performed using Fiji software.

### Cell morphology and sarcomere organization metrics

The sarcomere structure of micropatterned hiPSC-CMs or non-patterned was labeled with *α*-actinin antibody. Fluorescent images were acquired using a Cam mounted on a Leica TSC SP8 microscope, and confocal images were processed using Fiji. Morphological features and internal organization of sarcomeres were analyzed with an adapted MATLAB interface of SarcOmere Texture Analysis (SOTA) coded by Sutcliffe and al, (14). A total of 120 to 150 cells were analyzed per condition from three independent experiments.

### Mitochondrial morphology analysis

For mitochondrial network staining, live hiPSC-CMs from micropatterned or non-patterned condition (day 42) were labeled with 250 nM MitoTracker™ Orange CMTMRos (Thermo Fisher Scientific) in RPMI-1640/B27 plus insulin for at least 45 minutes at 37°C in a 5% CO2 incubator. Cells were then fixed and stained with *α*-actinin antibody as described above (Supplementary Table2). Fluorescent images were captured on a Cam mounted on a Leica SPE microscope with a 63x oil (n=1.3) objective. Z-stacks were deconvolved using Autoquant X software (MediaCybernetics, Bethesda, MD). Volumes of cells and individual mitochondrion were obtained with IMARIS software (Bitplane Company, Zurich, Switzerland), using Imaris MeasurementPro module. Fluorescence emitted from mitochondria was analyzed by the software allowing by surface segmentation methods, to identify individual objects corresponding to individual mitochondrion. Mitochondria segmented based on MitoTracker™ intensity were subject to single-cell quantification of surface area and volume in IMARIS and data were exported and categorized in Excel.

The alignment of mitochondrial network was visualized from MitoTracker™ Orange CMTMRos staining and confirmed using directionality analysis in Fiji software.

### Mitochondrial DNA quantification

Mitochondrial DNA content was quantified as described previously (26). Genomic and mitochondrial DNA was extracted with the NucleoSpin Tissue (Macherey-Nagel) from micropatterned and non-patterned hiPSC-CMs. The relative content of mtDNA was detected by quantitative real-time PCR using mitochondrially encoded tRNA leucine 1 (*MT-TL1*) primers and normalized *b-Actin*. All primers are listed in Supplementary Table3.

### Quantitative Real-Time PCR

Total RNA preparations were extracted with the NucleoSpin RNA Kit (MACHEREY-NAGEL) from the cells and quantified using a NanoDrop (Thermo Fisher Scientific). One microgram of total RNA was reverse transcribed to cDNA using the SuperScript™ IV VILO™ kit (Thermo Fisher Scientific) as per the manufacturer’s instructions. Quantitative real-time PCR was performed using amplified cDNA, gene-specific primers, and a SYBR Select Master Mix (Thermo Fisher Scientific) with the QuantStudio 3 Real-Time PCR System (Applied Biosystems). *RPL32* was used as the housekeeping gene. The relative gene expression was calculated by the 2^−ΔΔ*CT*^ method. All primers are listed in Supplementary Table3.

### Proteins extraction and Western Blotting

Total proteins were extracted from from micropatterned or non-patterned hiPSC-CMs (day 42) using RIPA buffer supplemented with a protease and phosphatase inhibitor cocktail (Sigma). Each sample was subjected to electrophoresis on 4-12% NuPAGE Bis-Tris gradient gels (Thermo Fisher Scientific), and proteins were transferred to Nitrocellulose membranes (Bio-rad). After blocking with 5% non-fat milk, the membranes were incubated overnight at 4°C with primary antibodies (Supplementary Table2). The next day membranes were incubated in secondary antibodies conjugated to horseradish peroxidase (Supplementary Table2) at room temperature for 1 hour. Finally, membranes were incubated with SuperSignal West Pico Plus Chemiluminescent Substrate (Thermo Fisher Scientific) for 5 minutes and luminescence was detected using the Amersham ImageQuant™ 800. Vinculin or Histone 3 were used as loading control protein. Densiometric analysis of the immunoblots was performed using ImageQuantTL Software (v.10.1.401, Cytiva).

### Mitochondrial and Glycolysis Bioenergetic Analysis

hiPSC-CMs from micropatterned or non-patterned conditions (day 42) were seeded into VN-coated Seahorse 96 assay plates (Seahorse Bioscience, Agilent Technologies France, Les Ulis, France) at a density of approximately 30,000 cells/well and allowed to grow for 4 days in RPMI-1640/B27 plus insulin medium. Mitochondrial oxidation and glycolytic function were evaluated by analysis of oxygen consumption rate (OCR, pmol/min/µg of protein) and extracellular acidification rate (ECAR, mpH/min/µg of protein) using the Seahorse XF96 extracellular flux analyser (Seahorse Bioscience), respectively. One hour prior, the culture medium was changed with the assay medium containing unbuffered DMEM supplemented with 10 mM Seahorse XF glucose solution (Agilent), 1 mM Seahorse XF pyruvate solution (Agilent) and 2 mM Seahorse XF glutamine (Agilent) for the mitochondrial oxidation assay or 4 mM L-glutamine for the glycolysis assay. Cells were maintained at 37°C without supplemental CO_2_ starting 1 hour before the assay. For the mitochondrial oxidation assay, OCRs were obtained from the slope change in oxygen concentration over time. After measurements of the baseline OCR, selective inhibitors were sequentially injected during the measurement (5 µM oligomycin, 1 µM FCCP and 1 µM antimycinA/rotenone, all from Sigma-Aldrich) following the manufacturer’s instructions. For normalization, cells were lysed in the XF96 plates using a protein lysis buffer (20 µL/well), and protein concentration was determined using the Bradford method.

For the glycolysis assay, Extracellular Acidification Rate (ECAR) was obtained from the slope of change in H^+^ concentration over time. After measurements of the baseline ECAR, selective inhibitors were sequentially injected during the measurements (10 mM glucose, 5 µM oligomycin, 50 mM 2-DG) following manufacturer’s instructions. All calculated values of ECAR parameters were normalized to quantified protein content.

### Mitochondrial oxygen consumption

Mitochondrial respiration was measured in digitonin-permeabilized hiPSC-CMs from micropatterned or non-patterned conditions (day 42) in an Oxygraph high-resolution respirometer chamber (Oroboros Oxygraph-2k, Innsbruck, Austria). Briefly, hiPSC-CMs were dissociated into single cells, counted, and washed with 1 mL of R solution (2.77 mM CaK_2_EGTA, 7.23 mM K_2_EGTA, 1.38 mM MgCl^2^, 3 mM KH_2_PO_4_ H_2_O, 20 mM imidazole, 20 mM taurine, 0.5 mM dithiothreitol, 90 mM K-methanesulfonate, 10 mM Na-methanesulfonate, 2 g/L FFA-free BSA; pH 7.1). After centrifugation, cells were transferred to 540 µL of R solution and inserted into chamber (O2k-sV-Module) to determine the rates of oxygen consumption at 37°C with continuous stirring as previously described (47). Reagents were added into the closed chamber using Hamilton syringes (Oroboros Instruments). For permeabilization of the cell membrane, 10 µg/ml of digitonin (Sigma) was added to the chamber at the start of the protocol until leak respiration (L) is stabilized. Mitochondrial electron transfer capacity (ETC) was then assessed following the sequential addition of 1 mM pyruvate/4 mM malate, 2 mM ADP, 10 mM succinate, 2 mM amytal, 10 µM oligomycin, and 0.5 mM TMPD combined with 2 mM ascorbate. The respiration rates were expressed in attomoles of O_2_ per second per cell.

### Duolink® Proximity Ligation In Situ Assay

Duolink® in situ proximity ligation assay (PLA) enables the detection, visualization, and quantification of protein-protein interactions (<40 nm) as individual fluorescent dots by microscopy. Cell fixation and permeabilization were performed as described for immunostaining. Blocking (1 hour at RT) and incubation with primary (Supplementary Table2) and secondary antibodies (1/5, 1 hour at 37°C) using anti-rabbit MINUS (DUO92005) and anti-mouse PLUS (DUO92001) kits, followed by ligation and amplification steps using the detection reagents orange kit (DUO92007), were performed according to the manufacturer’s protocol (Sigma-Aldrich). Coverslips were counter-stained with DAPI nuclear dye (1:10,000) and mounted with Dako Faramount Aqueous Mounting. Fluorescence was analysed on a Leica TSC SP8 microscope, 8-10 fields per condition for each experiment were acquired and the number of interactions per cell was then analyzed using Fiji software.

### Transcriptic analysis

We used hiPSC cells, non-patterned and micropatterned hiPSC-CMs at 42 days of differentiation to perform RNA-sequencing. Total RNA was extracted with a NucleoSpin RNA mini Kit (Macherey-Nagel), according to the manufacturer’s instructions. Library prep and RNA sequencing were conducted by Integragen as described below.

#### Quantification of gene expression

STAR was used to obtain the number of reads associated to each gene in the Gencode v31 annotation (restricted to protein-coding genes, antisense and lincRNAs). Raw counts for each sample were imported into R statistical software. The extracted count matrix was normalized for library size and coding length of genes to compute FPKM expression levels.

#### Unsupervised analysis

The Bioconductor edgeR package was used to import raw counts into R statistical software, and compute normalized log2 CPM (counts per millions of mapped reads) using the TMM (weighted trimmed mean of M-values) as normalization procedure. The normalized expression matrix from the 1000 most variant genes (based on standard deviation) was used to classify the samples according to their gene expression patterns using principal component analysis (PCA), hierarchical clustering and consensus clustering. PCA was performed by FactoMineR::PCA function with “ncp = 10, scale.unit = FALSE” parameters. Hierarchical clustering was performed by stats::hclust function (with Euclidean distance and ward.D method). Consensus clustering was performed by ConsensusClusterPlus::ConsensusClusterPlus function to examine the stability of the clusters. We established consensus partitions of the data set in K clusters (for K = 2, 3, …, 8), on the basis of 1,000 resampling iterations (80% of genes, 80% of sample) of hierarchical clustering, with Euclidean distance and ward.D method. Then, the cumulative distribution functions (CDFs) of the consensus matrices were used to determine the optimal number of clusters (K = 3 for instance), considering both the shape of the functions and the area under the CDF curves. tSNE analysis was performed with the Bioconductor Rtsne package applied to the PCA object (theta=0.0, perplexity=, max_iter=1000).

#### Differential expression analysis

The Bioconductor edgeR package was used to import raw counts into R statistical software. Differential expression analysis was performed using the Bioconductor limma package and the voom transformation. To improve the statistical power of the analysis, only genes expressed in at least one sample (FPKM >=) were considered. A qval threshold of <= 0.05 and a minimum fold change of 1.2 were used to define differentially expressed genes.

#### Differential pathway analysis

Differential pathway analysis was performed with the ReactomeGSA browser using the PADOG algorithm (Reactome version 82 on Nov, 2022). PADOG: Pathway Analysis with Down-weighting of Overlapping Genes (PADOG) computes a gene set score based on gene weights designed to emphasize the genes appearing in few gene sets, versus genes that appear in many gene sets (uniqueness of genes) (48). Differentially expressed pathways with an adjusted p-value below 0.05 were considered significantly regulated.

### Transmission Electron Microscopy

The cells were fixed by addition of glutaraldehyde (G5882, Sigma-Aldrich) in RPMI-1640-B27 with insulin media (final concentration 2.5% glutaraldehyde) for 30 minutes at room temperature, followed by a second fixation with 2.5% glutaraldehyde solution in 0.1 M sodium cacodylate buffer overnight at 4°C. Fixed cells were washed three times with PBS. A post-fixation was performed for 1 hour in a reduced osmium solution containing 1% osmium tetroxide, 1.5% potassium ferrocyanide in PBS 1x followed by incubatoin with a 1% thiocarbohydrazide solution in water for 20 minutes at room temperature, followed by 1% aqueous uranyl acetate at 4°C overnight. The samples were then subjected to *en* bloc Walton’s lead aspartate staining, and placed in a 60°C oven for 30 minutes. Then samples were dehydrated in graded series of ethanol for 10 minutes each. The samples were infiltrated with 50% Agar low viscosity resin (Agar Scientific Ltd) for 2 hours and 100% Agar low viscosity resin overnight. The resin was then changed by 100% Agar low viscosity for 1 hour, two times and the samples were further incubated for 1 hour before polymerization for 18 hours at 60°C. Sections with a thickness of 70 nm were cut using an EM UC6 ultramicrotome (Leica) and collected on formvar coated copper slot grids. Sections were observed at 120 kV with a Tecnail12 transmission electron microscope (Thermo Fisher Scientific) equipped with a 4Kx4K Oneview camera (Gatan)

### Statistical analysis

All numerical results are expressed as mean ± standard deviation (SD). Differences between experimental groups were analysed with the appropriate statistical tests, specified each time. P values < 0.05 were considered significant for all statistical tests. Statistical analyses were performed with GraphPad Prism software (V10.3.0).

## ACKNOWLEDGEMENTS

This work benefited from expertise and support from the iPSC technical facility at PARCC. We thank Camille Knops and Yunling Xu from the Flow Cytometry and the Microscopy platforms, respectively, from Université de Paris Cité, Paris Cardiovascular Research Center, Paris, France. We thank the ImagoSeine core facility of Institut Jacques Monod, member of France-BioImaging (ANR-10-INBS-04) and GIS-IBiSA. We would also like to thank Jérome Piquereau and Valérie Nicolas for helpful assistance with IMARIS analysis. We thank 4Dcell staff for the discussion about the cell repellent and design of the linear-shaped micropatterns. We thank Karima Ichegour for excellent administrative assistance.

## FUNDINGS

This work was supported by the French National Research Agency (NADHeart ANR-17-CE17-0015-02, NADINCARD ANR-22-CE92-0034-02), the Fondation pour la Recherche Médicale (EQU201903007852), and by a grant from the Leducq Foundation (18CVD05).

## DISCLOSURES

J.S.H. has received speaker, advisory board or consultancy fees from Alnylam, Amgen, Astra Zeneca, Bayer, Boerhinger Ingelheim, Novartis, Novo Nordisk, all unrelated to the present work. Other authors have no disclosures.

## Notes

### Competing Interest Statement

The authors have declared no competing interest.

