## Supplementary material for "Myocardial disarray drives metabolic inefficiency in human cardiomyocytes": Supp fig 1

### Supplementary figures

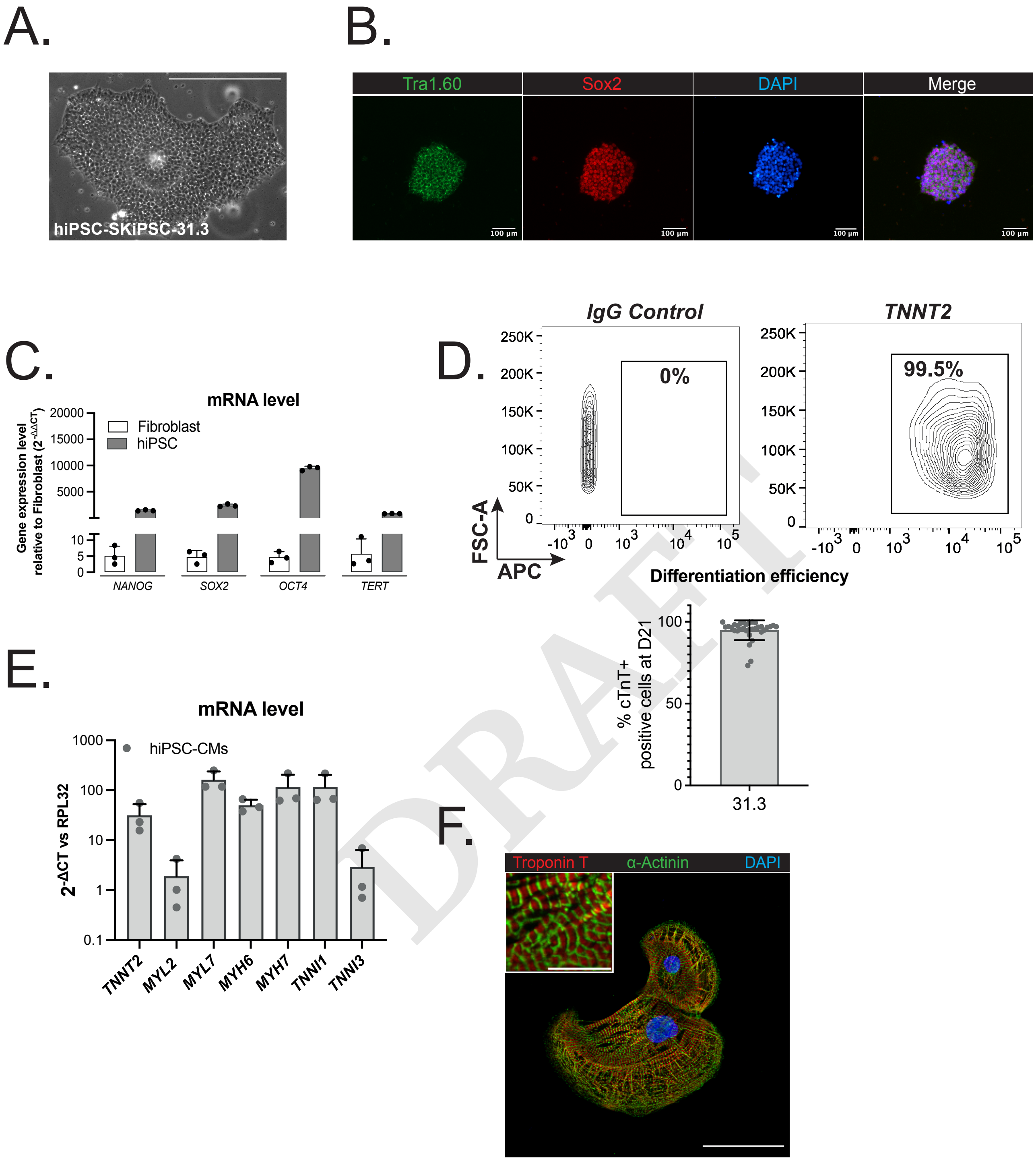


DRAFT

**Supplementary Figure 1. hiPSC and hiPSC-CMs characterization. A.** Representative phase-contrast image of the hiPSC cells. Scale bar 400 μm. **B.** Representative immunostaining in the hiPSC cells for Tra1.60 (green), and Sox2 (red). DAPI (blue) was used to stain nuclei. Scale bar 100 μm. **C.** Relative mRNA levels of pluripotency markers in hiPSC cells normalized to normal human dermal fibroblasts. **D.** Upper: Contour plot of hiPSC-CMs with TNNT2 staining analysed by FACS to calculate cardiac differentiation efficiency. 30,000 events recorded per group. Lower: Cardiac differentiation efficiency was calculated by the percentage of TNNT2$^{+}$ cells after 21 days of differentiation; n = 10 batches per group. **E.** Relative mRNA levels of cardiac markers hiPSC-CMs normalized to *RPL32*. N = 3 differentiations. **F.** Representative immunofluorescence images of hiPSC-CMs, for $\alpha$-sarcomeric actinin (green), and cardiac Troponin T2 (red) at day 30 after differentiation. DAPI (blue) was used to stain nuclei. The insert represents a zoomed view of sarcomeres. Global scale bar 50 μm, insert scale bar 10 μm.
