## Supplementary material for "Myocardial disarray drives metabolic inefficiency in human cardiomyocytes": Supp Fig 2

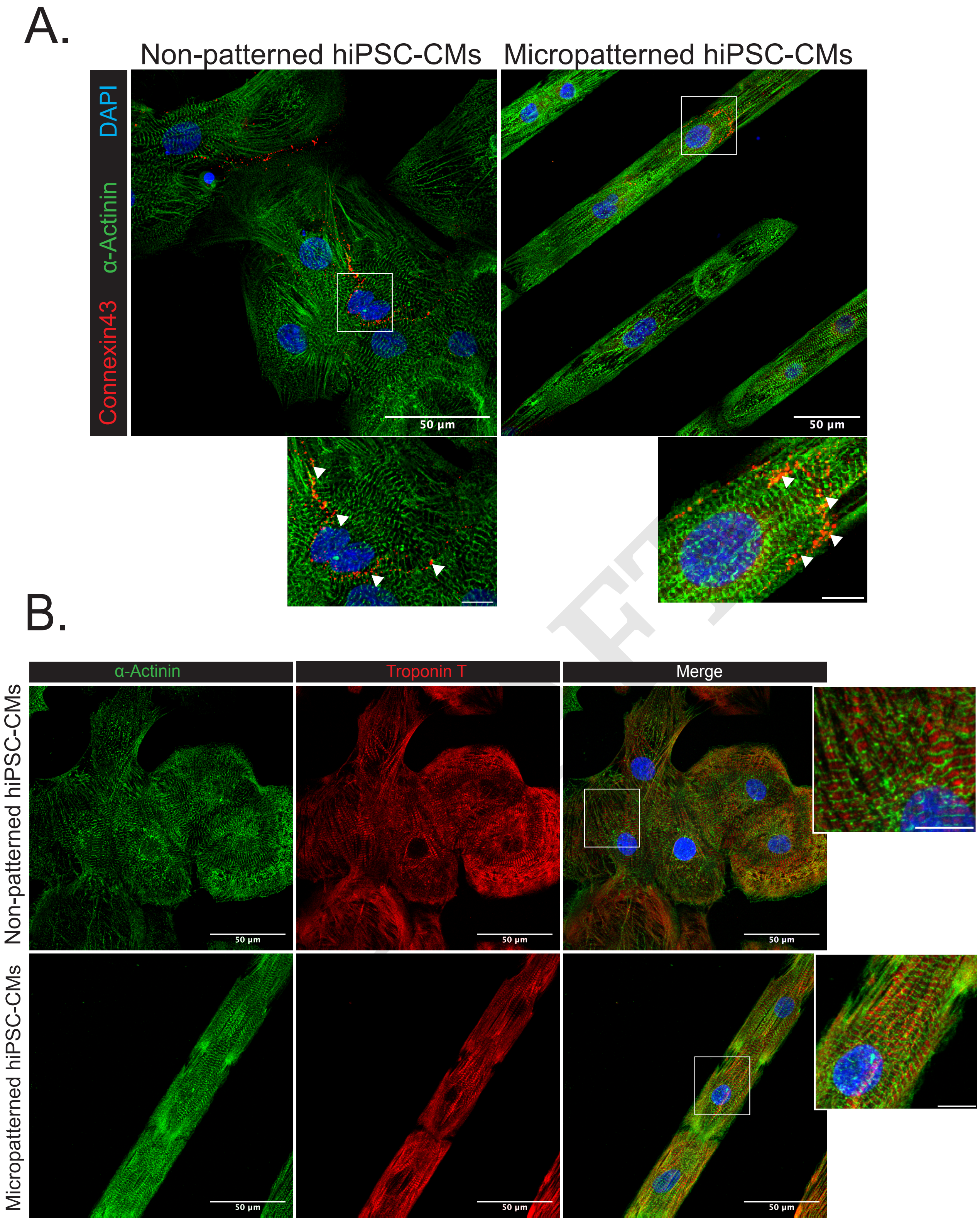


**Supplementary Figure 2. Comparison of sarcomeric structures in non-patterned and micropatterned hiPSC-CMs. A.** Non-patterned and micropatterned hiPSC-CMs stained with alpha-sarcomeric actinin (green) and connexin43 (red) 7 days after seeding. DAPI (blue) was used to stain nuclei. The insert represents a zoomed view of cell connection, scale bar 10 µm. **B.** Non-patterned and micropatterned hiPSC-CMs stained with alpha-sarcomeric actinin (green) and cardiac Troponin T2 (red) 7 days after seeding. DAPI (blue) was used to stain nuclei. The insert represents a zoomed view of sarcomeres, scale bar 10 µm.
