## Supplementary material for "Myocardial disarray drives metabolic inefficiency in human cardiomyocytes": Supp Fig 3

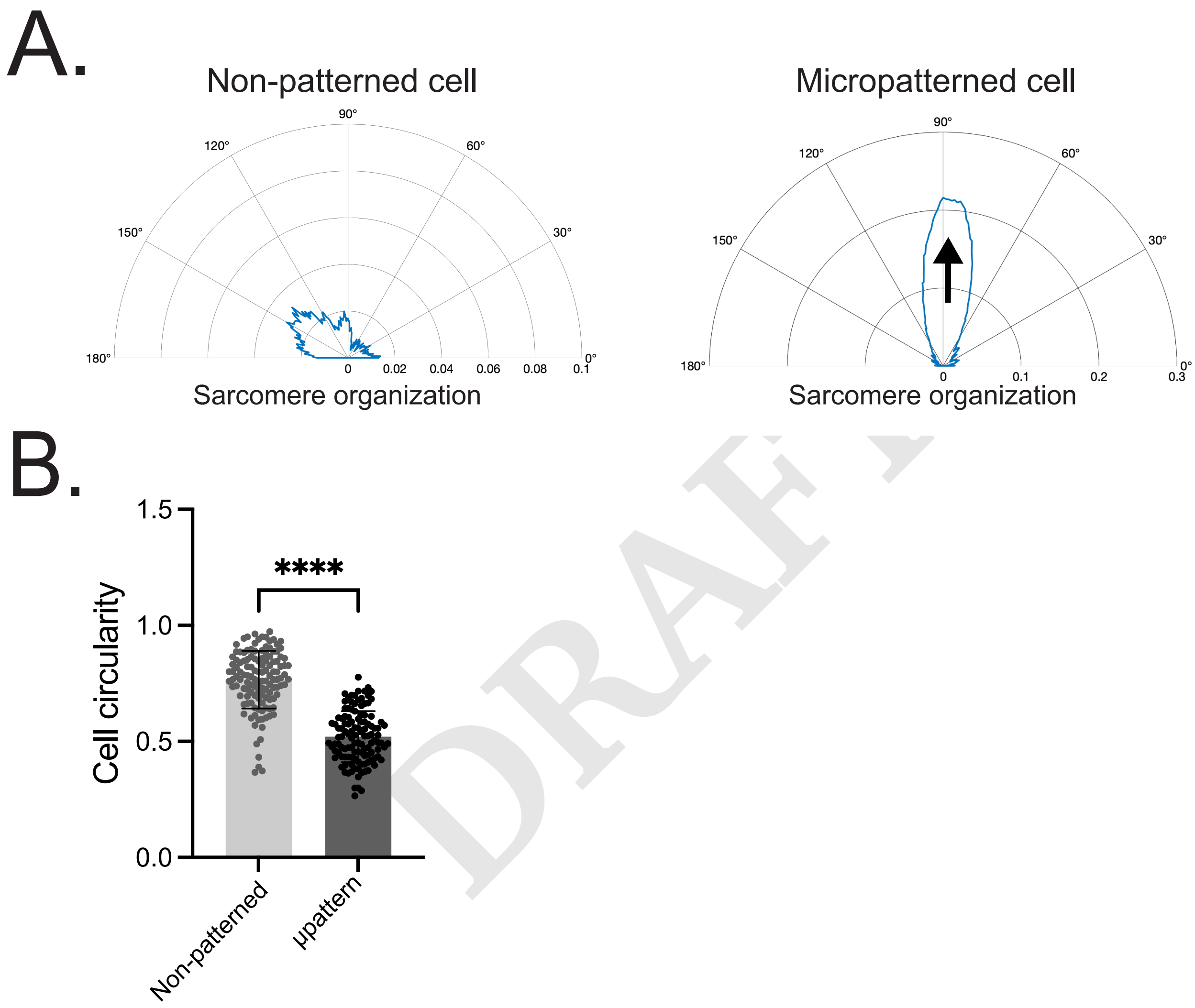


**Supplementary Figure 3. Automated measures of sarcomere organization. A.** Haralick correlation metric computed at multiple offset distances and angles to determine sarcomere organization and sarcomere length. Sarcomere organization assessed as a function of angle to assess the primary direction of sarcomere alignment. Arrow points in the direction of sarcomere alignment. **B.** Scatter plot shows cell circularity, measured with automated morphology measurements. n = 138 non-patterned cells, n = 120 micropatterned cells; N = 3-4 differentiations. Results, mean ± standard deviation. Statistical analyses: Unpaired t-test was used for B. **** p<0.0001 for the indicated comparisons.
