## Supplementary material for "Myocardial disarray drives metabolic inefficiency in human cardiomyocytes": Supp Fig 4

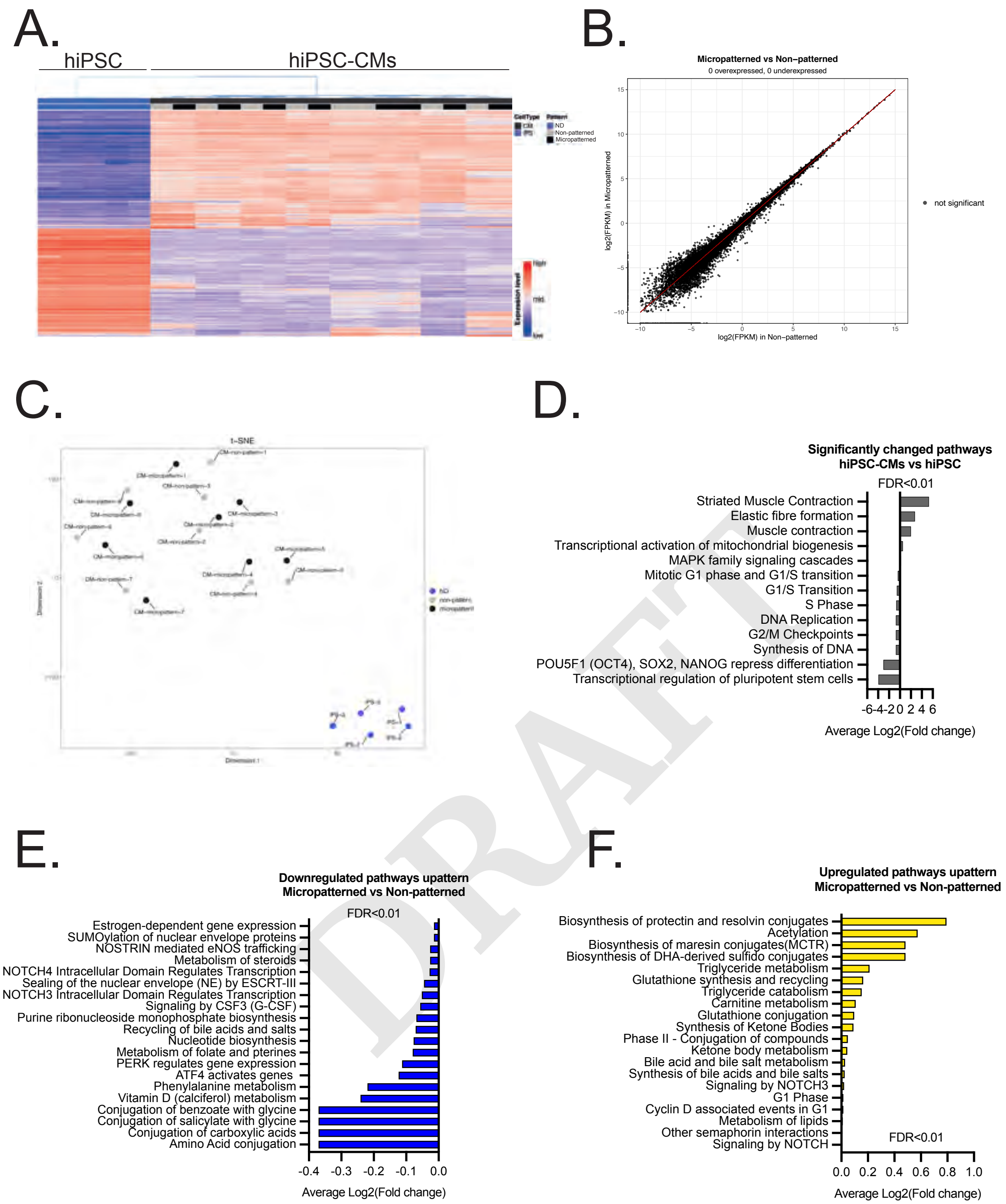


DRAFT

**Supplementary Figure 4. Significantly up- and down-regulated pathways in micropatterned hiPSC-CMs compared to non-patterned hiPSC-CMs A.** Heatmap of gene expression showing hierarchical clustering of hiPSC and hiPSC-CMs cells (micropatterned and non-patterned) detected by RNA-seq. N = 5-8 differentiations. **B.** Scatterplot depicting the Log2(FPKM) gene expression of micropatterned vs. non-patterned hiPSC-CMs. Black dots indicate no significant differential expression. **C.** tSNE plot of hiPSC (blue), hiPSC-CMs non-patterned (grey), and micropatterned (black) clusters. **D.** Representative significantly changed pathways (FDR>0.01) between hiPSC-CM and hiPSC as found with ReactomeGSA pathway analysis. Data presented as the average Log2(Fold change). **E.** All significantly downregulated pathways (FDR<0.01) in micropatterned vs non-patterned hiPSC-CMS detected with ReactomeGSA. Data presented as the average Log2(Fold change). **F.** All significantly upregulated pathways (FDR<0.01) in micropatterned vs non-patterned hiPSC-CMS detected with ReactomeGSA. Data presented as the average Log2(Fold change).
