## Supplementary material for "Myocardial disarray drives metabolic inefficiency in human cardiomyocytes": Supp Fig 5

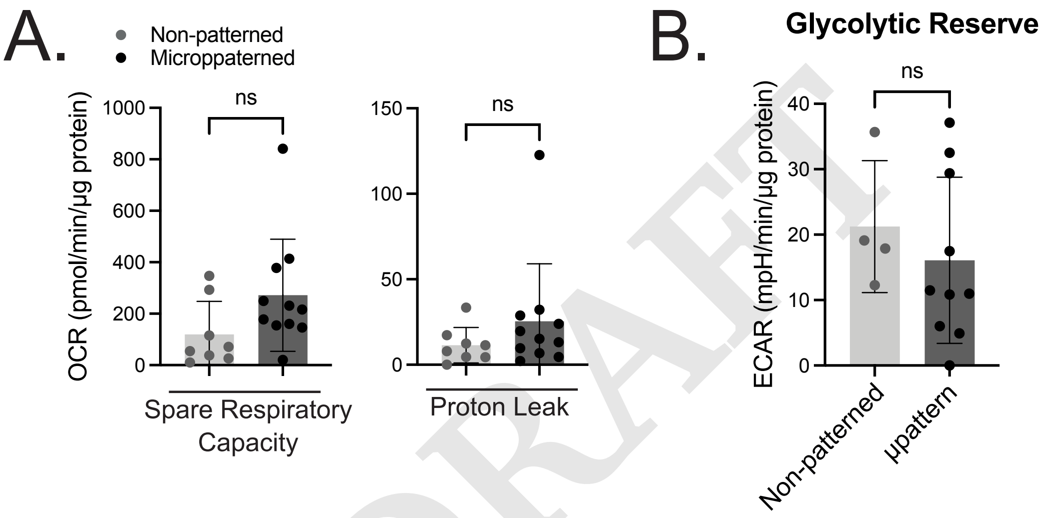


DRAFT

**Supplementary Figure 5. Additional results of mitochondrial respiration and glycolytic function in micropatterned hiPSC-CMs compared to non-patterned hiPSC-CMs. A.** Scatter plots show spare respiratory capacity, and proton leak. N = 8-11 differentiations. **B.** Quantification of glycolytic reserve in non-patterned and micropatterned group. N = 4-12 differentiations. Results, mean $\pm$ standard deviation. Statistical analyses: Mann-Whitney was used for A, and B. * p<0.05, ns: not significant, for the indicated comparisons.
