## Supplementary material for "Myocardial disarray drives metabolic inefficiency in human cardiomyocytes": Supp Fig 6

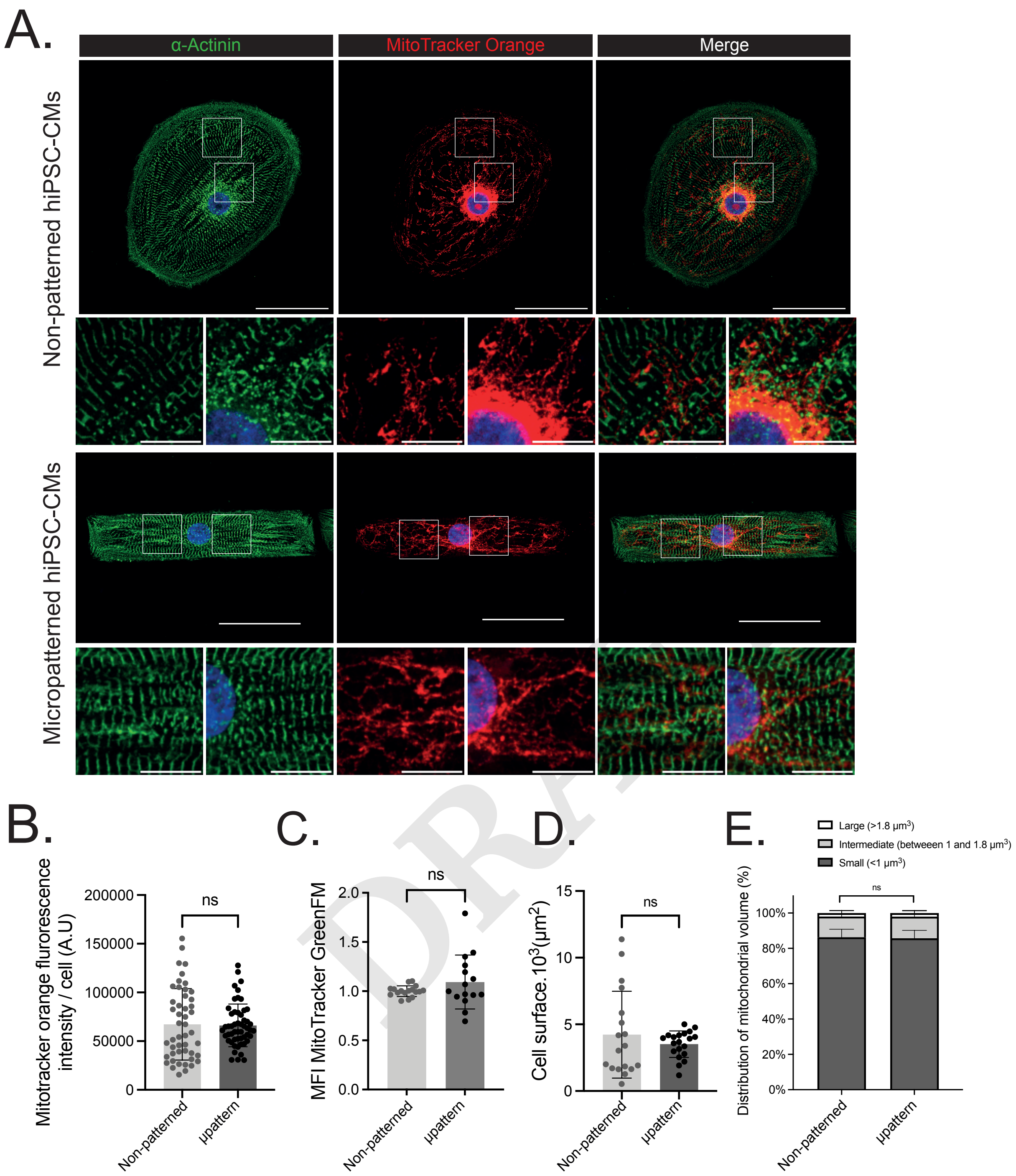


**Supplementary Figure 6. Mitochondrial network analysis in micropatterned and non-patterned hiPSC-CMs. A.** Non-patterned and micropatterned hiPSC-CMs stained with MitoTracker™ Orange 7 days after seeding. DAPI (blue) was used to stain nuclei. The insert represents a zoomed view of sarcomeres and mitochondrial networks. Global scale bar 50 µm, insert scale bar 10 µm. **B.** Quantitative analysis of MitoTracker™ Orange fluorescence intensity measured with Fiji software. n = 48 non-patterned cells, n = 53 micropatterned cells; N = 3-4 differentiations. **C.** Flow cytometry analysis of MitoTracker™ Green in non-patterned and micropatterned hiPSC-CMs. N = 6 differentiations. **D.** Scatter plot shows cell surface measured with Fiji software. n = 18 non-patterned cells, n = 20 micropatterned cells; N = 3 differentiations. **E.** Distribution of individual mitochondrial volume between small (<1 $\mu m^3$), intermediate (between 1 and 1.8 $\mu m^3$) or large (>1.8 $\mu m^3$) mitochondria evaluated by IMARIS. Results, mean $\pm$ standard deviation. Statistical analyses: Mann-Whitney was used for B through D, Kruskal-Wallis test followed post-hoc Dunn's multiple comparisons test for E. ns: not significant for the indicated comparisons.
