## Supplementary material for "Myocardial disarray drives metabolic inefficiency in human cardiomyocytes": Supp Fig 7

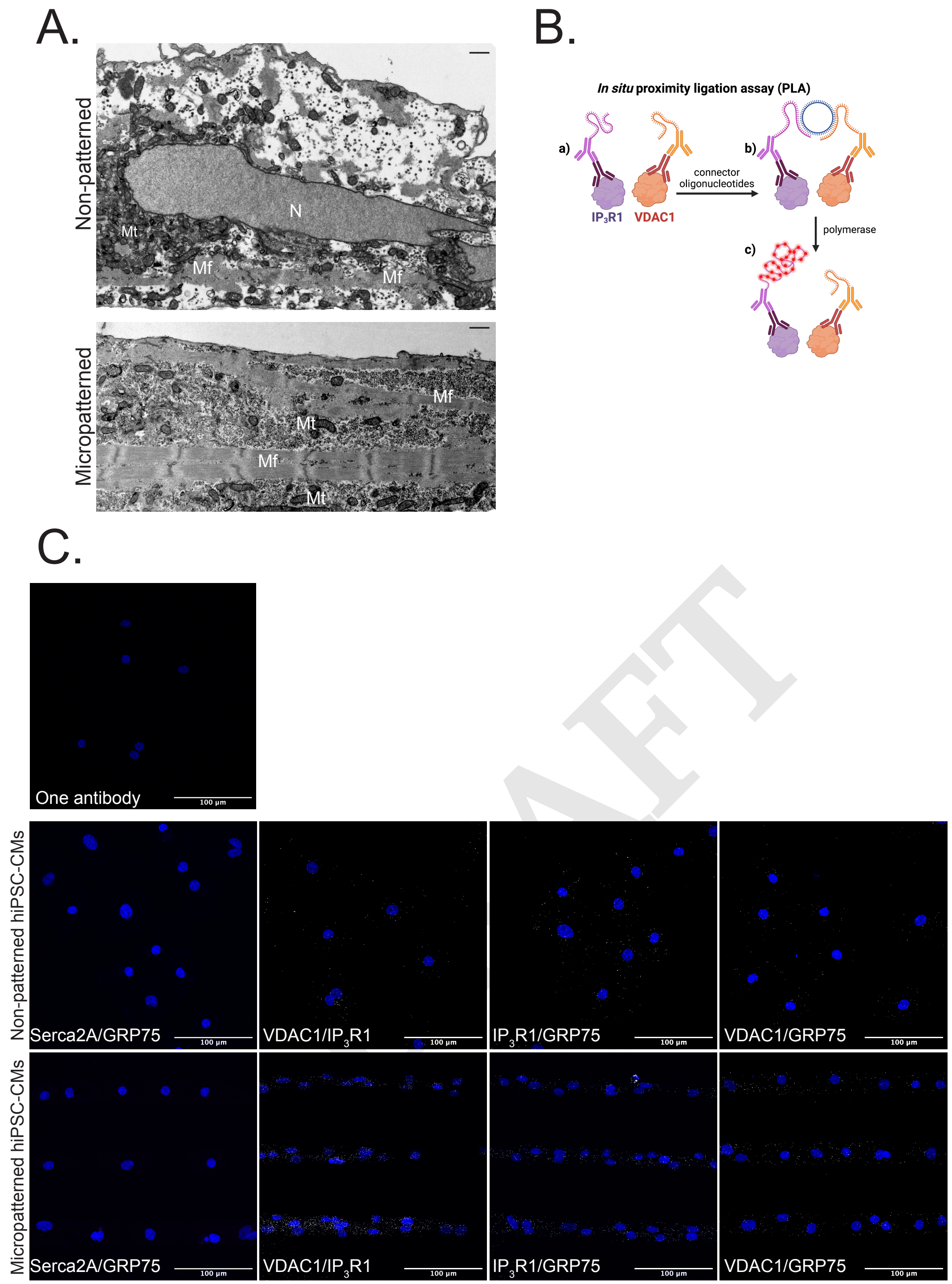


**Supplementary Figure 7. Ultra structural characteristics of micropatterned and non-patterned hiPSC-CMs. A.** Representative TEM images of the cellular structure of non-patterned (top) and micropatterned (bottom) hiPSC-CMS in transverse cross-section; scale = 1 µm. **B.** Schematic representation of in situ PLA steps for visualization of VDAC1/IP$_3$R1 interactions: a) fixed and permeabilized cultured hiPSC-CMs are incubated with two different primary antibodies, one directed against the mitochondrial channel VDAC1 and another one against the IP$_3$R1 channel in the SR, b) circularization and ligation of connector oligonucleotides of secondary antibodies when proteins are less than 40 nm away, c) rolling circle amplification with polymerase and detection of the product with fluorescent probes. **C.** Representative images of in situ PLA analysis in hiPSC-CMs cells. Nuclei appear in blue and interactions between two targeted proteins are depicted in whites. IP$_3$R1 interacts with both VDAC1 and GRP75. As negative controls, we demonstrate that GRP75 do not interacts with Serca2a (another SR membrane protein).
