## Supplementary material for "Myocardial disarray drives metabolic inefficiency in human cardiomyocytes": Supp Tables 1 and 2

### Supplementary tables

**Table 1.** 16 entities identified in the carnitine metabolism pathway from the REACTOME Pathway Analysis. Up/down expressions correspond to changes of plus or minus 30% respectively.

| Gene | UniPro Id | logFC | Interpretation |
|---|---|---|---|
| ACACA | Q13085 | -0,17461268 | down |
| ACACB | O00763 | 0,09898097 | |
| CPT1A | P50416 | 0,243642567 | up |
| CPT1B | Q92523 | 0,222166436 | up |
| CPT2 | P23786 | 0,085895949 | |
| MID1IP1 | Q9NPA3 | -0,213965579 | down |
| PPARD | Q03181 | -0,152778742 | down |
| PRKAA2 | P54646 | 0,052515816 | |
| PRKAB2 | O43741 | 0,002802349 | |
| PRKAG2 | Q9UGJ0 | 0,091991203 | |
| RGMB | O00863 | -0,128303368 | |
| RXRA | P19793 | 0,055295189 | |
| SLC22A4 | O76082 | -0,178795184 | down |
| SLC22A5 | O76082 | -0,162658222 | down |
| SLC25A20 | O43772 | 0,102333938 | |
| THRSP | Q92748 | 1,252323355 | up |

**Table 2.** Antibodies used for Immunofluorescence (IF), Duolink PLA, Flow cytometry (FC) and Western blot (WB).

| Antibodies | Manufacturer | Assay | Cat. No. | Dilution |
|---|---|---|---|---|
| cTnT, APC, REAfinity | Miltenyi Biotech | FC | 130-120-403 | 1:100 |
| REA control human IgG1, APC, REAfinity | Miltenyi Biotech | FC | 130-120-709 | 1:100 |
| cTnT | Abcam | IF | ab45932 | 1:500 |
| $\alpha$-actinin | Sigma-Aldrich | IF | A7811 | 1:1000 |
| Connexin 43 | Thermo Fisher Scientific | IF | 71-0700 | 1:200 |
| Tra1.60 | Millipore | IF | MAB4304 | 1:100 |
| SOX2 | Millipore | IF | AB5603 | 1:200 |
| VDAC1 | Abcam | PLA/WB | ab14734 | 1:200/1:1000 |
| VDAC | Thermo Fisher Scientific | WB | PA1-954A | 1:1000 |
| IP$_3$R1 | Millipore | PLA | 07-1213 | 1:100 |
| IP$_3$R1 | Cell Signaling | WB | 8568 | 1:1000 |
| GRP75 | Cell Signaling | PLA/WB | 3593 | 1:100/1:1000 |
| GRP75 | Santa Cruz | PLA | sc-133137 | 1:100 |
| OPA1 | BD Biosciences | WB | 612607 | 1:1000 |
| TOM40 | Proteintech Group | WB | 18409-1-AP | 1:1000 |
| Total OXPHOS Cocktail | Abcam | WB | ab110411 | 1:1000 |
| SDHA | Invitrogen | WB | 459200 | 1:1000 |
| Histone 3 | Abcam | WB | ab1791 | 1:1000 |
| Vinculin | Sigma-Aldrich | WB | V9131 | 1:5000 |
| AMPK$\alpha$ | Cell Signaling | WB | 5831 | 1:1000 |
| ACC | Cell Signaling | WB | 3676 | 1:1000 |
| p-AMPK(Thr172) | Cell Signaling | WB | 2535 | 1:1000 |
| p-ACC(Ser72) | Cell Signaling | WB | 3661 | 1:1000 |
| Rabbit HRP Conjugated | Jackson ImmunoResearch | WB | 711-035-152 | 1:10,000 |
| Mouse HRP Conjugated | Jackson ImmunoResearch | WB | 715-035-150 | 1:10,000 |
| Goat anti-Rabbit IgG (H+L) HRP | Euromedex | WB | BE-A120-101P | 1:10,000 |
| Goat anti-Mouse IgG (H+L) HRP | Euromedex | WB | BE-A90-116P | 1:10,000 |
| Goat anti-Rabbit IgG (H+L), Alexa Fluor® 546 | Thermo Fisher Scientific | IF | A11010 | 1:1000 |
| Goat anti-Mouse IgG (H+L), Alexa Fluor® 488 | Thermo Fisher Scientific | IF | A10680 | 1:1000 |

DRAFT
