## Supplementary material for "Myocardial disarray drives metabolic inefficiency in human cardiomyocytes": Supp Table 3

**Table 3.** List of primers.

| Target | Forward primer | Reverse primer | Product |
|---|---|---|---|
| RPL32 | AGTTCCTGGTCCACAACGTC | GTGACTCTGATGGCCAGTTG | 142 |
| NANOG | CTCCAACATCCTGAACCTCAGC | CGTCACACCATTGCTATTCTTCG | 115 |
| SOX2 | GCCCTGCAGTACAACTCCAT | GACTTGACCACCGAACCCAT | 128 |
| OCT4 (POU5F1) | AATTTGTTCCTGCAGTGCCC | CTCTCGTTGTGCATAGTCGC | 150 |
| hTERT | CAAGCTGTTTGCGGGGATTC | AGTTCACCACTGTCTTCCGC | 167 |
| TNNT2 | CAGGATCAACGATAACCAGAAAGTC | GTGAAGGAGGCCAGGCTCTA | 87 |
| MYL2 | GCAGGCGGAGAGGTTTTC | AGTTGCCAGTCACGTCAGG | 74 |
| MYL7 | CCGTCTTCCTCACGCTCTT | TGAACTCATCCTTGTTCACCAC | 120 |
| MYH6 | TCTCCGACAACGCCTATCAGTAC | GTCACCTATGGCTGCAATGCT | 140 |
| MYH7 | GGCAAGACAGTGACCGTGAAG | CGTAGCGATCCTTGAGGTTGTA | 133 |
| TNNI1 | TCCGTGGGAAGTTCAAGCG | GACTTGGCGGCATCAAACATC | 238 |
| TNNI3 | CCAACTACCGCGCTTATGC | CTCGCTCCAGCTCTTGCTTT | 120 |
| MT-TL1 | CGAAAGGACAAGAGAAATAGAG | GAACAAGGTTTTAAGTCTTACGCA | 134 |
| $\beta$-Actin | GTGACGTTGACATCCGTAAAGA | CCTCACCAAGCTAAGGATGC | 250 |

DRAFT
